# Extended DNA binding interface beyond the canonical SAP domain contributes to SDE2 function at DNA replication forks

**DOI:** 10.1101/2022.05.09.490802

**Authors:** Alexandra S. Weinheimer, YiTing Paung, Julie Rageul, Arafat Khan, Brian Ho, Michael Tong, Sébastien Alphonse, Markus A. Seeliger, Hyungjin Kim

**Author notes:** These authors contributed equally (Alexandra S. Weinheimer, YiTing Paung).

## Abstract

Elevated DNA replication stress causes instability of the DNA replication fork and DNA mutations, which underlies tumorigenesis. The DNA replication stress regulator SDE2 binds to TIMELESS (TIM) of the fork protection complex (FPC) and enhances its stability, thereby supporting replisome activity at DNA replication forks. Here, we structurally and functionally characterize a new conserved DNA binding motif related to SAP (<u>S</u>AF-A/B, <u>A</u>cinus, <u>P</u>IAS) in human SDE2 and establish its preference for single-stranded DNA (ssDNA). The nuclear magnetic resonance solution structure of SDE2^SAP^ reveals a helix-extended loop-helix core aligned parallel to each other, consistent with known canonical SAP folds. Notably, its DNA interaction extends beyond the core SAP domain and is augmented by two lysine residues in the C-terminal tail, which is uniquely positioned adjacent to SAP and conserved in the pre-mRNA splicing factor SF3A3. Mutation in the SAP domain with extended C-terminus not only disrupts ssDNA binding but also impairs TIM localization at replication forks, thus inhibiting efficient fork progression. Together, our study establishes SDE2^SAP^ as an essential element for SDE2 to exert its role in preserving replication fork integrity via FPC regulation and highlights the structural diversity of the DNA-protein interactions achieved by a specialized DNA binding motif.

## INTRODUCTION

DNA replication is one of the most fundamental processes for the survival of organisms, and therefore needs to be completed efficiently and accurately. DNA replication is coordinated by the replication machinery, or the replisome, at DNA replication forks, where a Cdc45-MCM-GINS (CMG) helicase complex unwinds duplex DNA, while proliferating cell nuclear antigen (PCNA) acts as a processivity factor to guide the synthesis of leading and lagging strands by replicative polymerases ε and δ (1). Disruption of DNA replication fork progression that stalls or blocks polymerization processes causes DNA replication stress, which activates a range of checkpoints and repair pathways that are tightly regulated (2). Importantly, multiple DNA-protein interactions occur at sites of DNA replication and damage, and many replication regulators, DNA repair enzymes, and signal transducers harbor specific DNA binding motifs to coordinate controlled DNA transactions necessary for replication fork integrity (3, 4). Defects in the DNA replication stress response cause single-stranded DNA (ssDNA) accumulation and strand breakage, culminating in chromosome aberrations and genome instability that are often observed in cancer (5). Persistent DNA replication stress is a key hallmark of cancer, and the DNA replication vulnerability of cancer cells is being exploited as a new therapeutic target (6).

Replication fork stability is reinforced by the fork protection complex (FPC), a heterodimer of TIMELESS (TIM)-TIPIN along with ancillary proteins AND1/Ctf4 and CLASPIN/Mrc1, which together act as a scaffold to couple the polymerase-helicase activities thereby supporting seamless replication fork progression (7–11). Recent cryo-EM structural analyses of the replisome complex in human and yeast revealed positioning of the TIM-TIPIN heterodimer ahead of CMG to grip duplex DNA and promote strand separation (12, 13). AND1, existing as a trimer in the complex, helps stabilize replication forks by coupling the primase Pol α to helicase function and promotes sister chromatid cohesion establishment (14, 15). The TIM-TIPIN heterodimer also transmits the ssDNA-RPA signal at stalled forks to initiate the replication checkpoint, in which CLASPIN directly interacts with CHK1 to facilitate ATR-dependent CHK1 phosphorylation (16–18). Thus, the FPC serves as an essential platform to coordinate DNA replication and the DNA damage response.

We recently identified Human SDE2 (silencing-defective 2, C1orf55) as a new regulatory component of the FPC at active and stalled replication forks (19). Originally discovered as a PCNA-associated protein at sites of DNA replication, SDE2 directly interacts with TIM to promote its stability and localization to replication forks (19–21). SDE2 deficiency phenocopies the loss of TIM, and the SDE2-TIM interaction is required for ensuring efficient fork progression and protecting stalled forks from nucleolytic degradation (19). Given its structural role without any obvious enzymatic activity, SDE2 may contribute to the tethering of TIM to replication forks, thereby stabilizing the FPC in the replisome. Intriguingly, we previously showed that chromatin-dependent degradation of N-terminally cleaved SDE2 is necessary for propagating the signals of the replication stress response at ssDNA-accumulated stalled forks under UVC damage, indicating that the DNA binding property of SDE2 may determine the levels of replisome-associated SDE2 and its function at replication forks (22). However, exactly how SDE2 interacts with DNA to exert its role in replication fork integrity remains unclear.

Structural elements within SDE2 provide an important clue for SDE2 regulation. SDE2 contains a SAP domain at its C-terminus, named after its original identification in the DNA scaffold attachment factors (<u>S</u>AF)-A and SAF-B, <u>A</u>cinus, and protein inhibitor of activated STAT1 (<u>P</u>IAS) (23). The SAP domain is an evolutionarily conserved helix-loop-helix motif, which exhibits a bipartite distribution of hydrophobic and polar amino acids with an invariant glycine in the loop. This short ∼35-residue motif is considered a DNA binding motif predominantly present in proteins involved in DNA damage signaling and repair, including Ku70, PARP1 (from *A. thaliana*), RAD18, and SLX4, suggesting that its function may be specialized for genome maintenance. Intriguingly, the SAP domain of SDE2 is present in all major metazoans but is absent in yeast, indicating that SDE2 may have acquired additional functions in the DNA damage response throughout its evolution, besides its role in mRNA splicing and telomere maintenance characterized in *S. pombe* Sde2 (24, 25). Despite the discovery of the SAP domain two decades ago, very few studies have been conducted on it, and there is still little known about its biological functions associated with genome maintenance.

In this study, we describe a new conserved DNA binding motif related to SAP in human SDE2 and determine its role in regulating SDE2 function within the FPC at DNA replication forks. The SDE2^SAP^ preferentially binds to ssDNA, and its canonical SAP fold is extended to the C-terminal tail that is uniquely present in SDE2, both of which contribute to ssDNA interaction. This unique configuration is conserved in the pre-mRNA splicing factor subunit SF3A3, defining an extended SAP domain. We further show that SDE2^SAP^ is required for localizing TIM at replication forks and ensuring replication fork progression in cells. Together, our study establishes that the extended SAP domain constitutes an essential element necessary for the biological activity of SDE2 in replication fork stabilization and provides new insight into the versatility of the DNA-protein interactions that contribute to genome maintenance and suppression of tumorigenesis.

## RESULTS

### The SAP domain is required for SDE2 to bind DNA and localize at replication forks

Human SDE2 contains three evolutionarily conserved domains in its 451 amino acid (aa) polypeptide (**Fig. 1A**). The N-terminal 77 aa constitutes a ubiquitin-like (UBL) domain, which is cleaved at its C-terminal di-glycine motif by deubiquitinating enzyme activity, releasing a processed C-terminal SDE2 (SDE2^Ct^) at replication forks (20). The coiled-coil SDE2 domain is required for the interaction of SDE2 with TIM, which promotes stable association of the FPC at active replication forks (19). Under DNA damage, regulated chromatin-associated degradation of SDE2 promotes the RPA activation necessary for the stress response at stalled replication forks (22). Given the importance of SDE2 function at replication forks, we sought to determine whether its putative DNA binding domain of SDE2, the SAP domain (SDE2^SAP^), mediates SDE2 binding to DNA, and how the DNA binding property of SDE2 controls its function at replication forks. As with other proteins, the SAP domain of SDE2 is predicted to exhibit a bipartite distribution of hydrophobic and polar residues separated by a linker loop that contains a glycine residue (**Fig. 1A**). These key residues are all conserved in metazoans (**Fig. S1A**). Subcellular fractionation revealed that deletion of the SAP domain abolishes the localization of SDE2 to the chromatin-enriched fraction (**Fig. 1, B and C**). Additionally, fluorescence loss in photobleaching (FLIP) showed that intranuclear mobility of the GFP-tagged SAP mutant is increased in comparison to wild-type, indicating that the SAP domain is required for the stable association of SDE2 with chromatin (**Fig. S1B**). To confirm its specific localization at active replication forks, we employed a proximity ligation assay (PLA) modified for detecting proteins at nascent DNA labeled by 5-Ethynyl-2’-deoxyuridine (EdU) (i.e. *in situ* analysis of protein interactions at DNA replication forks, or SIRF) (26). As reported previously (20), cells expressing wild-type SDE2 exhibited distinct SDE2-EdU PLA foci, while the SAP mutant-expressing cells showed a significant decrease in PLA foci and numbers within, indicating that the SAP domain is necessary for localizing SDE2 at sites of DNA replication (**Fig 1, D and E, and Fig. S1C**). Moreover, the SAP mutant purified from U2OS cells failed to be pulled down by biotinylated ssDNA, indicating that the SAP domain is required for DNA binding (**Fig. 1, F and G**). To further confirm the DNA binding property of the SAP domain *in vitro*, we purified recombinant full-length SDE2 wild-type or the SAP deletion mutant from *E. coli* (**Fig. S1D**). Electrophoretic mobility shift assays (EMSAs) with fluorescently labeled DNA oligonucleotides revealed that wild-type SDE2 binds to DNA, predominantly to ssDNA and splayed-arm DNA (saDNA), which mimics a replication fork, and the SAP mutation abolished its binding to DNA (**Fig. 1, H and I**). Addition of SDE2 or Flag antibodies, but not a tubulin antibody, resulted in a supershift with ssDNA but not with double-stranded DNA (dsDNA), confirming the specificity of DNA binding in EMSA (**Fig. 1J** **and Fig. S1E**). Together, these data support that SDE2^SAP^ is a bona fide DNA binding motif and suggest that this binding property is necessary for SDE2 localization at replication forks.

**Figure 1.**
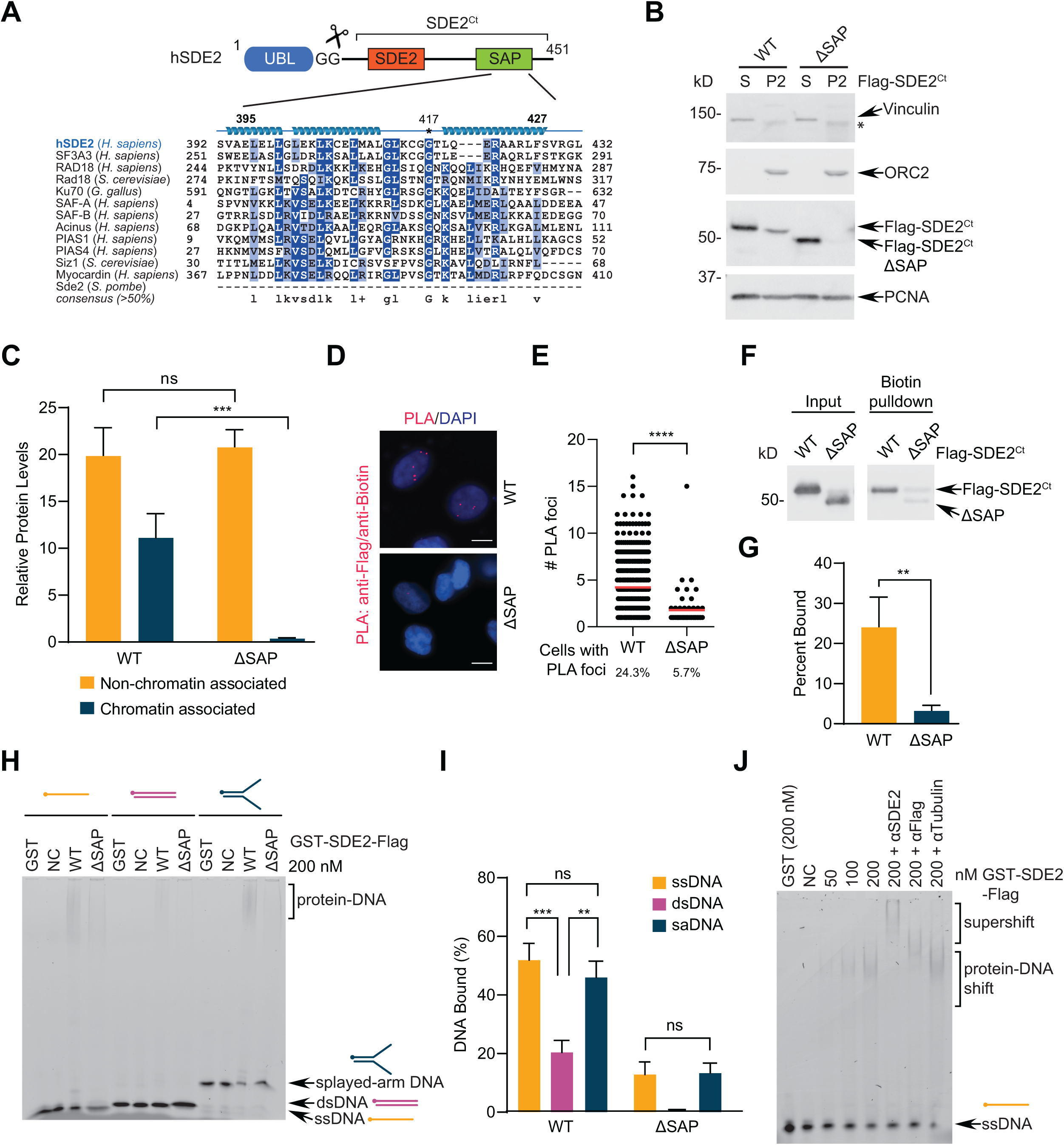
The SAP domain of SDE2 is required for DNA binding. **(A)** Schematic of SDE2 domain structures and sequence alignment of SDE2^SAP^ with various SAP-domain containing proteins. **(B)** U2OS cells transfected with either Flag-SDE2^Ct^ WT or ΔSAP (Δ395-451) were fractionated to separate cytosolic and chromatin-associated protein pools and analyzed by Western blotting. S indicates cytosolic proteins, P2 indicates acid-soluble chromatin-bound proteins. Asterisk (*) indicates non-specific bands. **(C)** Quantification of (B) from more than three independent experiments; error bar: SEM, \*\*\**p*<0.001, two-tailed Mann-Whitney test. ns, not significant. **(D)** U2OS cells were transfected with either Flag-SDE2 WT or ΔSAP, and proximity to the replication fork was analyzed using the SIRF proximity ligation assay. Scale bar: 10 μm **(E)** The average number of PLA foci per cell. Quantification of three independent experiments (>400 cells per condition); red bar: mean, \*\*\*\**p*<0.0001, two-tailed Mann-Whitney test. Percentage underneath graph indicates percent of total cells positive of PLA signal. **(F)** Proteins purified from U2OS cells transiently expressing Flag-SDE2 WT or ΔSAP were pulled down with biotinylated 80-mer ssDNA oligo and streptavidin magnetic beads, and analyzed by Western blotting. **(G)** Quantification of (F) from four independent experiments; error bar: SEM, \*\**p*<0.01, two-way ANOVA. **(H)** Electrophoretic mobility shift assay (EMSA) of purified GST-SDE2-Flag WT or ΔSAP (Δ395-427) incubated with 6-carboxyfluorescein (6-FAM)-labeled 60-mer ssDNA, dsDNA, or splayed-arm DNA. DNA: 5 nM, protein: 200 nM. NC = negative control, no protein. **(I)** Quantification of (H) from four independent experiments; error bar: SEM, \*\**p*<0.01, \*\*\**p*<0.001, two-way ANOVA. **(J)** EMSA of purified GST-SDE2-Flag incubated with 60-mer ssDNA. Where indicated, antibodies were added at a 1:2 ratio for a supershift EMSA. DNA: 50 nM, protein: 50-200 nM.

### SDE2^SAP^ preferentially binds single-stranded DNA

We noticed that SDE2 binds dsDNA nearly 3-fold less than the other DNA structures while performing the EMSAs and decided to examine this property in further detail. While wild-type SDE2 readily binds ssDNA and saDNA, even at low concentrations, SDE2 only binds dsDNA when provided in excess (**Fig. 2, A and B**). SDE2 binds to saDNA and ssDNA-dsDNA junction DNA (jnDNA) similarly to ssDNA, indicating that SDE2 recognizes ssDNA-containing structures (**Fig. S2A**). Biotinylated DNA pull-down using purified recombinant proteins further showed that SDE2 preferentially associates with ssDNA (**Fig. 2, C and D**). Similarly, the GST-tagged SAP domain (GST-SDE2^SAP^) alone bound ssDNA significantly better than dsDNA (**Fig. 2, E and F**). Due to the small size of GST-SDE2^SAP^, disappearance of free probe was used to determine fractions bound indirectly. As a control, an SDS-containing denaturing gel dissociated the SDE2-DNA complex, while no visible decrease in free probes was observed, arguing against the possibility that disappearance of the probe is due to nuclease contamination (**Fig. S2B**). Additionally, we employed fluorescence anisotropy to quantify preferential DNA binding affinity of the SAP domain. GST-SDE2^SAP^ caused a noticeable increase in anisotropy with increasing concentration, while binding to dsDNA causes an anisotropy increase barely above the background. From this data, the dissociation constant (*K*_d_) of SAP binding to ssDNA was calculated to be ∼1.77 μM. A similar result was observed from the fluorescence anisotropy with full-length SDE2 (**Fig. S2C**). Overall, our results suggest that ssDNA is a preferable substrate of SDE2^SAP^. Notably, a similar preference to ssDNA over dsDNA has been previously reported in RAD18^SAP^ (27).

**Figure 2.**
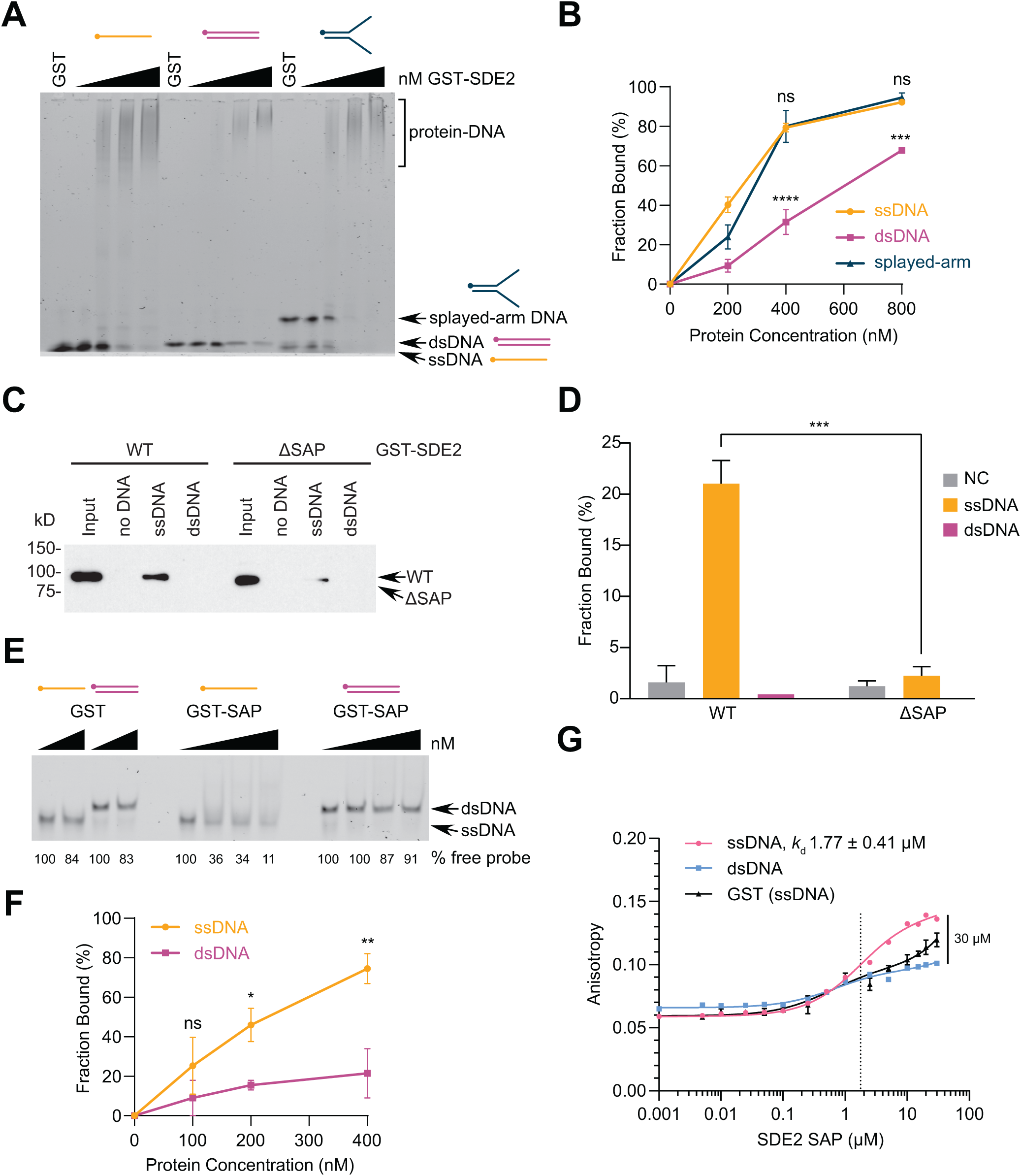
SAP preferentially binds single-stranded DNA. **(A)** EMSA of purified GST-SDE2-Flag incubated with 60-mer ssDNA, dsDNA, or splayed-arm DNA. **(B)** Quantification of (A) from three independent experiments; error bar: SEM, \*\*\**p*<0.001, \*\*\*\**p*<0.0001, two-way ANOVA. **(C)** Purified GST-SDE2 WT or ΔSAP (Δ395-427) was incubated with biotinylated 80-mer ssDNA oligo for biotin pull-down and Western blotting. **(D)** Quantification of (C) from two independent experiments; error bar: SEM, \*\*\**p*<0.001, two-way ANOVA. **(E)** EMSA of purified, recombinant GST-SDE2^SAP^ (aa. 350-451) incubated with 60-mer ssDNA or dsDNA. **(F)** Quantification of (E) from three independent experiments; error bar: SEM, \**p*<0.05, \*\**p*<0.01, two-way ANOVA. **(G)** Increasing amounts of GST-SDE2^SAP^ was incubated with FAM-labeled 60-mer ssDNA vs. dsDNA and analyzed using fluorescence anisotropy. The SAP domain binds ssDNA with a dissociation constant (*K*_d_) of 1.77 ± 0.41 μM. Data fit to a one-site, total binding saturation curve. Because a plateau was not reached, a *K*_d_ for dsDNA binding was not calculated.

### Residues in the loop region of the SAP domain are important for DNA binding

Previous analyses based on the predicted surface of RAD18^SAP^ and Ku70^SAP^ implicated residues that lie in the loop region close to the tips of the two helices in DNA binding (27, 28). Specifically, it was shown that mutations of conserved histidine, glycine, and leucine at the start of the loop or of glycine and lysine at the end of the loop have an adverse effect on the affinity of RAD18^SAP^ to ssDNA (27). To test whether the analogous residues in SDE2^SAP^ are important for DNA binding, we made point mutations of the corresponding residues, excluding histidine which is not considered conserved in our alignment (**Fig. S3A**). Mutation of these four conserved loop-region residues (G412A/L413E/G417A/L419E; GLGL) resulted in a significant decrease in the ability of SDE2 to localize to the chromatin (**Fig S3, B and C**). This result indicates that the mode of DNA binding may be generally preserved in the SAP domain despite some variations in the conserved loop-helix junctions from multiple SAP-containing proteins.

### The C-terminal tail of SDE2 contributes to SDE2^SAP^-dependent DNA binding

To determine the full extent of the DNA binding interface within SDE2^SAP^, we monitored DNA binding of purified SDE2^SAP^ (from aa 380 to aa 451; total 77 aa including five remnant residues upon GST cleavage) using solution nuclear magnetic resonance (NMR) by watching for chemical shift perturbation upon successive titration with ssDNA (**Fig. 3A**). ^1^H-^15^N HSQC spectra of unbound SDE2^SAP^ showed distinct and well-dispersed chemical shifts, indicative of a well-folded protein. Upon successive titration with a 16-mer ssDNA oligonucleotide, several cross-peaks, including those from C415, G417, T418, K444, and K447, were perturbed substantially upon DNA binding, reflecting potential DNA interactions (**Fig. 3, B and C**). Some minor changes were observed for K405, L413, K414, G416, E435, F442, and L446, possibly indicating minor conformational change upon DNA binding (for full spectra assignment, see **Fig. S4A**). The largest chemical shift perturbations were concentrated in the extended loop region, including the invariant G417, the conserved residue responsible for RAD18^SAP^ binding to DNA. This is consistent with our mutagenesis studies based on RAD18^SAP^, supporting a general mechanism for SAP domain interaction with DNA. Unexpectedly, there were also prominent chemical shift perturbations for residues K444 and K447, both located in the short tail that follows the helices in SDE2^SAP^ at its C-terminus (**Fig. 3, B and C**). This 24-amino acid stretch constitutes a C-terminal tail (SDE2^CTT^) that is evolutionally conserved but is uniquely present in SDE2 (**Fig. 3D**). For instance, Ku70, which harbors its SAP domain at the end of its C-terminus, does not exhibit this tail motif (28). Intriguingly, two lysine residues and others in SDE2^CTT^ as well as SDE2^SAP^ are well conserved within the middle region of the splicing factor SF3A3, a subunit of the pre-mRNA splicing complex, implicating that these two domains may have been evolutionarily selected as one functional unit that contributes to DNA binding (24).

**Figure 3.**
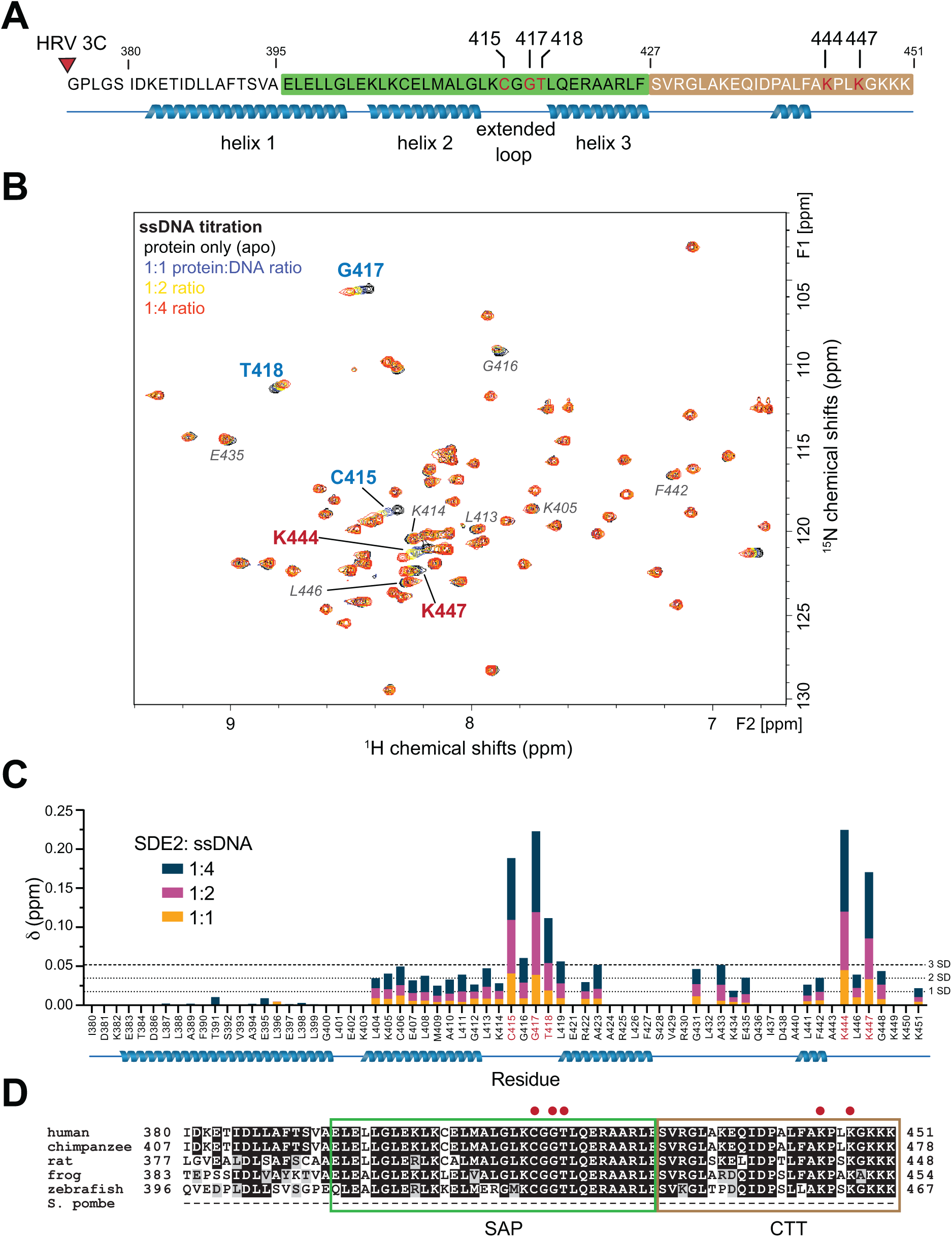
Analysis of DNA-interacting residues of SDE2. **(A)** Schematic showing a peptide used for the heteronuclear single quantum coherence (HSQC) experiment. The peptide was expressed with a GST tag, which was cleaved using HRV 3C protease (red triangle), leaving behind the SAP domain and C-terminal tail of SDE2. **(B)** Two-dimensional ^1^H-^15^N HSQC NMR spectra of purified SDE2^SAP^ (aa. 380-451) titrated with increasing ratios of 16-mer ssDNA. Overlaid spectra show perturbation of several cross-peaks upon addition of ssDNA, indicating interaction between SDE2^SAP^ and 16-mer ssDNA. Major shifts, indicative of DNA binding, are in bold. Minor perturbations, which are less likely to be from direct DNA interaction, are in italics. **(C)** Graph showing the extent of perturbation for each individual residue backbone. The areas showing the largest degree of movement are the extended loop and the C-terminus. 1 SD (standard deviation) = 0.0174; 2 SD = 0.0348; 3 SD = 0.0522. **(D)** Sequence alignment of SAP and C-terminal tail (CTT) of SDE2 from multiple species. Red dots indicate the binding residues identified from HSQC.

### The extended SDE2 SAP domain forms a helix-extended loop-helix motif connected to a unique C-terminal tail

Intrigued by the unique DNA binding mode of SDE2^SAP^, we solved the solution structure of residues 380-451, which encompasses the entire extended SAP domain, by NMR (statistics in **Table 1**). The core of polypeptides forms a well-defined structure that is composed of three primary helices, where helices 2 and 3 constitute a classic SAP motif (**Fig. 4A** **and Fig. S4B**). Like other SAP domains, SDE2^SAP^ folds into a helix-extended loop-helix motif, in which the two middle helices from the SAP core are connected to an interhelical loop and oriented nearly parallel to each other. The major DNA binding residues are concentrated to this interhelical loop, exposed away from the conserved hydrophobic core of the protein. Superposition of the SAP fold with a known SAP domain derived from Ku70 (29) and the AlphaFold-predicted model of SDE2^SAP^ revealed a high structural similarity, including orientation of helices 2 and 3, and formation of a hydrophobic core that stabilizes the structure (**Fig. 4B**). The core SAP fold is followed by a C-terminal tail (SDE2^CTT^), which is not present in any other previously characterized SAP domains but situated close to the interhelical loop of SDE2 in its three-dimensional structure, creating a cleft surrounded by the DNA interacting residues from the SAP core and the CTT (**Fig. 4C**). A template-based modeling predicts that ssDNA is able to fit in with this cleft, likely making contacts with both the loop and the tail, implicating to a potential mechanism of ssDNA-specific DNA interaction (**Fig. 4D**). Positively charged residues, lysine 444 and 447, in the tail are expected to augment the relatively weak interactions from the SAP core. Together, our structural analysis of SDE2^SAP^ suggests a unique mode of ssDNA binding mediated by the concerted interactions supported by the core SAP and CTT domains.

**Figure 4.**
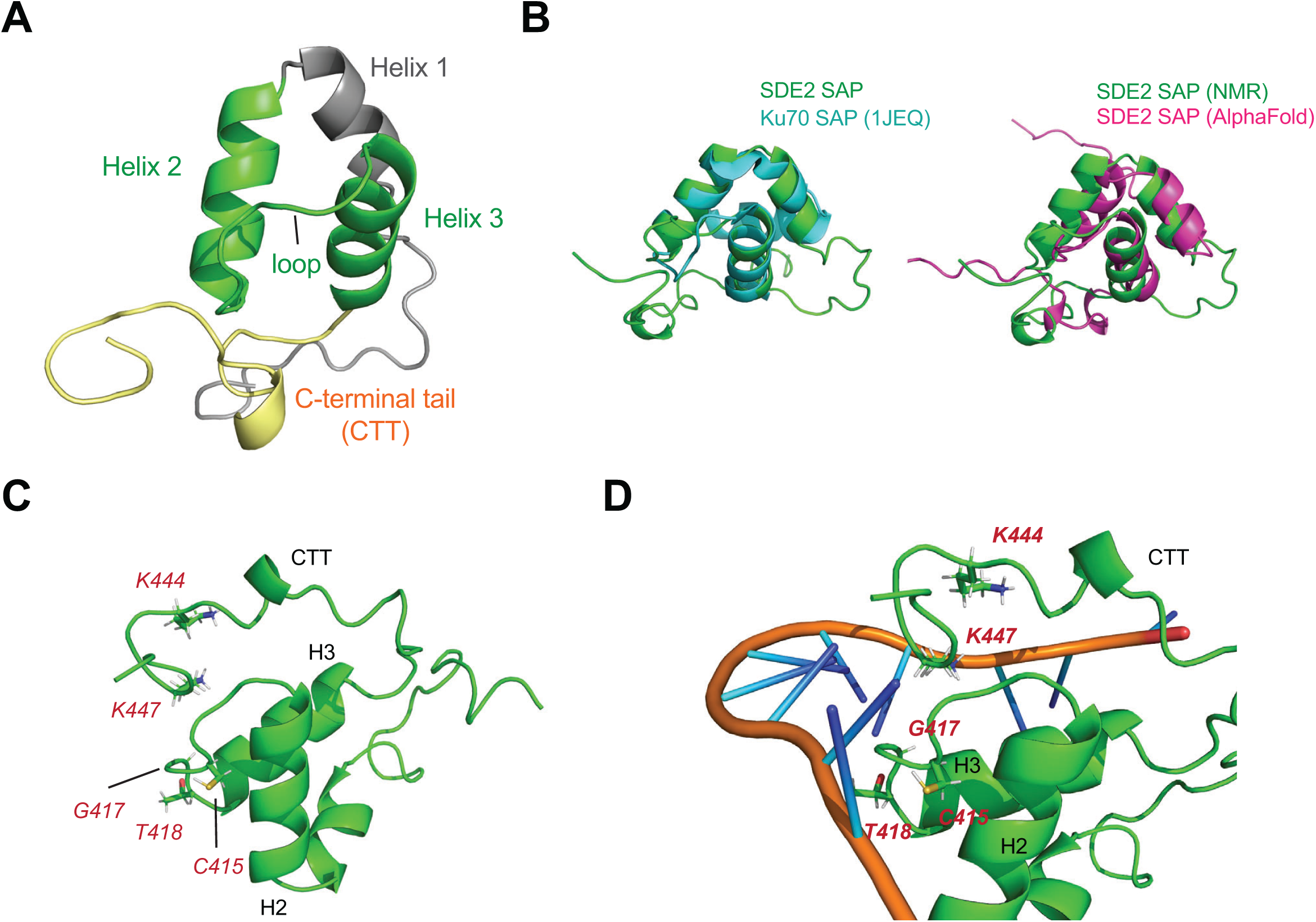
The NMR solution structure of the extended SDE2 SAP domain. **(A)** The backbone, displayed as a ribbon diagram, of the solution structure of the SAP+CTT motif of SDE2. The two helices and connecting residues that constitute the core SAP motif are in green. **(B)** Overlay of SDE2^SAP^ with a known SAP domain structure derived from Ku70 (PDB: 1JEQ) and the AlphaFold-predicted SDE2^SAP^. Despite high structural similarity in the SAP core, SDE2 harbors the CTT, which is not present in the SAP at the C-terminus of Ku70. **(C)** The locations of the most-perturbed residues during ssDNA binding, highlighted in red. The perturbed residues are grouped into a cleft formed by the interhelical loop within the core SAP and near the end of the CTT. **(D)** Docking simulation of ssDNA binding to the core SAP and the CTT using HDOCK Server. ssDNA structure was obtained from PDB ID: 5ZG9.

**Table 1.** The restraints and structural statistics for the 20 lowest energy conformers of SDE2^SAP^

| Distance Constraints |  |
| --- | --- |
| Intra-residue | 327 |
| Inter-residue | 214 |
| Short ($\vert i-j\vert \leq 1$) | 119 |
| Medium ($\vert i-j\vert < 5$) | 66 |
| Long ($\vert i-j\vert \geq 5$) | 29 |
| Hydrogen | 22 |
| Ambiguous | 104 |
| Residual dipolar restraints | 28 |
| Dihedral angle restraints | 58 |
| Violations |  |
| Distance violations ($> 0.5 \text{ \AA}$) | 0 |
| Angle violations ($> 5^\circ$) | 0 |
| Procheck Ramachandran Statistics (%) |  |
| Most favored regions | 86.4 |
| Additional allowed regions | 13.8 |
| Generously allowed regions | 1.5 |
| Disallowed region | 0 |
| Structural Statistics |  |
| NOE rmsd | 0.034 |
| Dihedral rmsd | 1.31 |
| RDC Q scores | 0.188 |
| RDC R correlation | 0.979 |
| RMSD |  |
| Heavy atoms ($\text{\AA}$) | $1.35 \pm 0.21 \text{ \AA}$ |
| Backbone atoms ($\text{\AA}$) | $0.67 \pm 0.2 \text{ \AA}$ |

### The extended SDE2^SAP+CTT^ mediates DNA interactions

Our NMR analysis suggests that the C-terminal tail may help SDE2 achieve higher affinity during DNA binding. In order to determine whether SDE2^CTT^ is an element required for efficient DNA binding by SDE2, we generated a series of N-terminal Flag-tagged SDE2 wild-type and mutants (**Fig. 5A**). Subcellular fractionation showed that deletion of either SDE2^SAP^ or SDE2^CTT^ significantly impairs the ability of SDE2 to localize at chromatin, and deletion of both domains completely abolished it, indicating that SDE2^SAP^ or SDE2^CTT^ cooperatively binds to DNA (**Fig. 5, B and C**). To further substantiate our findings, we determined the DNA binding affinity of SDE2 wild-type and mutants using EMSA. *In vitro* analysis using recombinant SDE2 proteins showed a similar result, where deletion of both domains is required for completely abrogation of SDE2 binding to ssDNA (**Fig. 5, D and E, and Fig. S5A**). Importantly, point mutations of the key residues in the SDE2^SDE2+CTT^ domains, identified with the HSQC study, abolished its binding to ssDNA, confirming the requirement of the CTT for DNA interaction (**Fig. 5F** **and Fig. S5B**). Disruption of two lysine residues in the tail was sufficient to significantly reduce the binding to ssDNA, further highlighting the importance of SDE2^CTT^ in mediating DNA interactions (**Fig. 5G**). These results indicate that both SDE2^SAP^ and SDE2^CTT^ function as independent yet compulsory elements in DNA binding and define the extended SDE2^SAP+CTT^ as an unconventional SAP responsible for the interaction of SDE2 with DNA.

**Figure 5.**
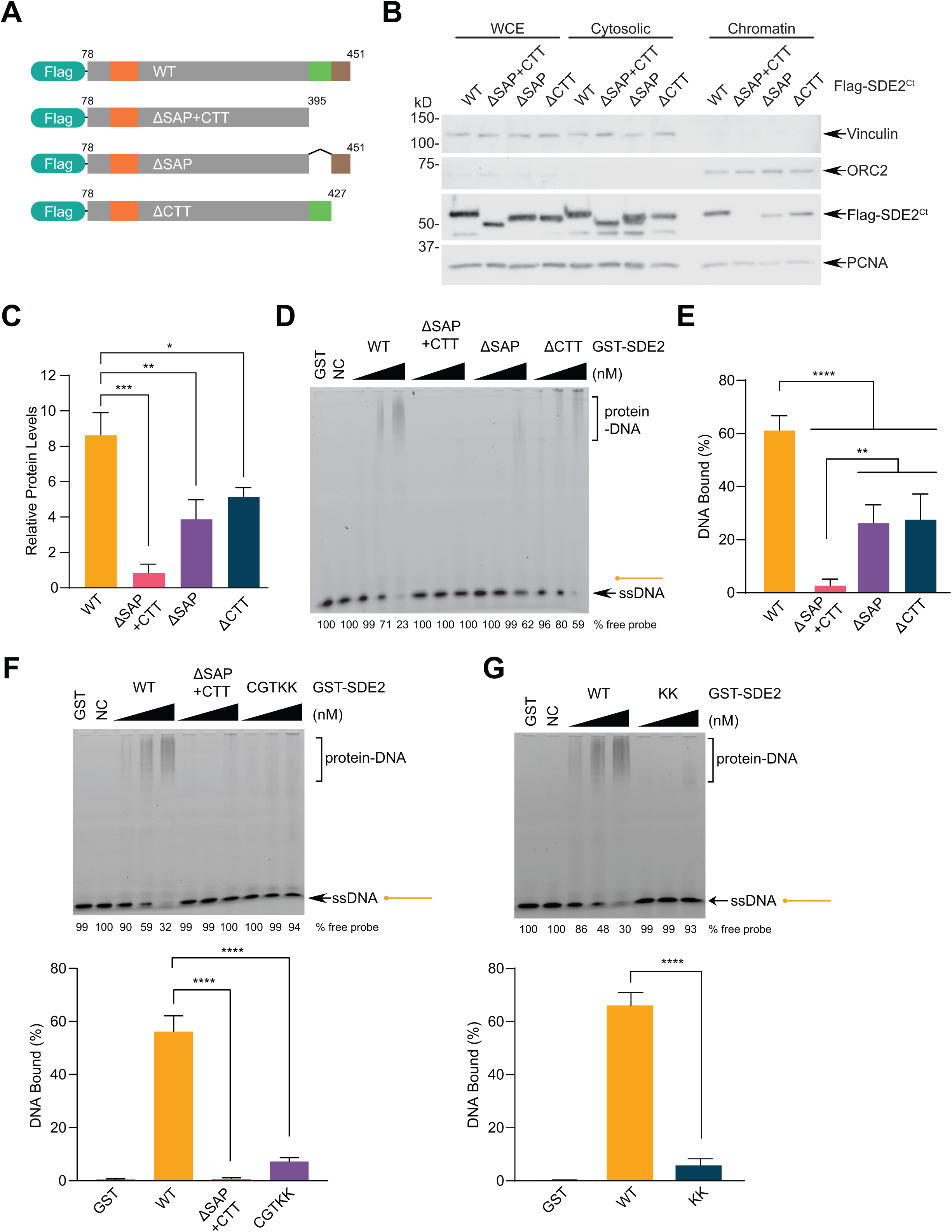
The C-terminal tail (CTT) of SDE2 contributes to ssDNA binding. **(A)** Schematic showing the Flag-SDE2^Ct^ deletion mutants used to determine the contribution of the SAP domain versus the C-terminal tail in DNA binding of SDE2. **(B)** U2OS cells were transfected with Flag-SDE2^Ct^ WT, ΔSAP+CTT, ΔSAP, or ΔCTT, fractionated to separate cytosolic and chromatin-associated protein pools, and analyzed by Western blotting. **(C)** Quantification of chromatin-associated Flag-SDE2^Ct^ from four independent experiments; error bar: SEM, \**p*<0.05, \*\**p*<0.01, \*\*\**p*<0.001, two-way ANOVA. **(D)** EMSA of purified GST-SDE2 (WT, ΔSAP+CTT, ΔSAP, or ΔCTT) incubated with 60-mer ssDNA. DNA: 25 nM, protein: 50, 100, 200 nM. NC = negative control, no protein. **(E)** Quantification of DNA-bound GST-SDE2 at 200 nM from four independent experiments; error bar: SEM, \*\**p*<0.01, \*\*\*\**p*<0.0001, two-way ANOVA. **(F, G)** EMSA of purified GST-SDE2 (WT, ΔSAP+CTT, CGTKK, KK) incubated with 60-mer ssDNA. DNA: 20 nM, protein: 50, 100, 200 nM. CGTKK: C415A/G417S/T418V/K444A/K447A. KK: K444A/K447A. For each panel, quantification of DNA-bound GST-SDE2 at 200 nM from four independent experiments is shown; error=SEM, \*\*\*\**p*<0.0001, two-way ANOVA.

### SDE2 and SF3A3 both contain the unique extended SAP+CTT motif

The identification of a new, extended SAP domain prompted us to investigate whether its unique mode of DNA binding, encompassing both the core SDE2^SAP^ and SDE2^CTT^, is generally applicable to other SAP-containing proteins. To this end, we examined SF3A3, a subunit of the pre-mRNA splicing complex that harbors a similar SAP+CTT domain structure (24, 30). This distinctive feature is not present in other SAP-containing proteins to the best of our knowledge, and both SDE2 and SF3A3 exhibit a remarkable degree of sequence conservation in the extended SAP region, including residues identified in our SDE2^SAP^ NMR ^1^H-^15^N HSQC analysis, compared to other SAPs, including RAD18^SAP^ (**Fig. 6A**). Notably, biotin pull-down showed a strong interaction of purified GST-SF3A3 to ssDNA (**Fig. 6B**). Importantly, deletion of either the SAP or CTT region of Flag-tagged SF3A3 partially abrogated the association of SF3A3 to chromatin, and deletion of both prevented its chromatin binding almost completely (**Fig. 6, C and D**). These results substantiate our finding in SDE2 that both the core SAP fold and the adjacent tail region contribute to DNA binding, and that this unique mode of DNA binding via the extended SAP can be applicable to other proteins with a similar domain configuration. It is interesting to note that Prp9, the *S. cerevisiae* homolog of SF3A3, harbors a zinc finger (ZnF) motif where the CTT is found in SF3A3, while mutations to the conserved cysteine and histidine residues render the motif nonfunctional (**Fig. S6A**) (31, 32). Therefore, although the ZnF motif, known to bind RNA in pre-mRNA splicing factors, was lost in the metazoan SF3A3^SAP+CTT^, it may have been evolutionarily selected for an acquired ability to recognize ssDNA, which was then transferred to SDE2 alongside the core SAP as a single unit.

**Figure 6.**
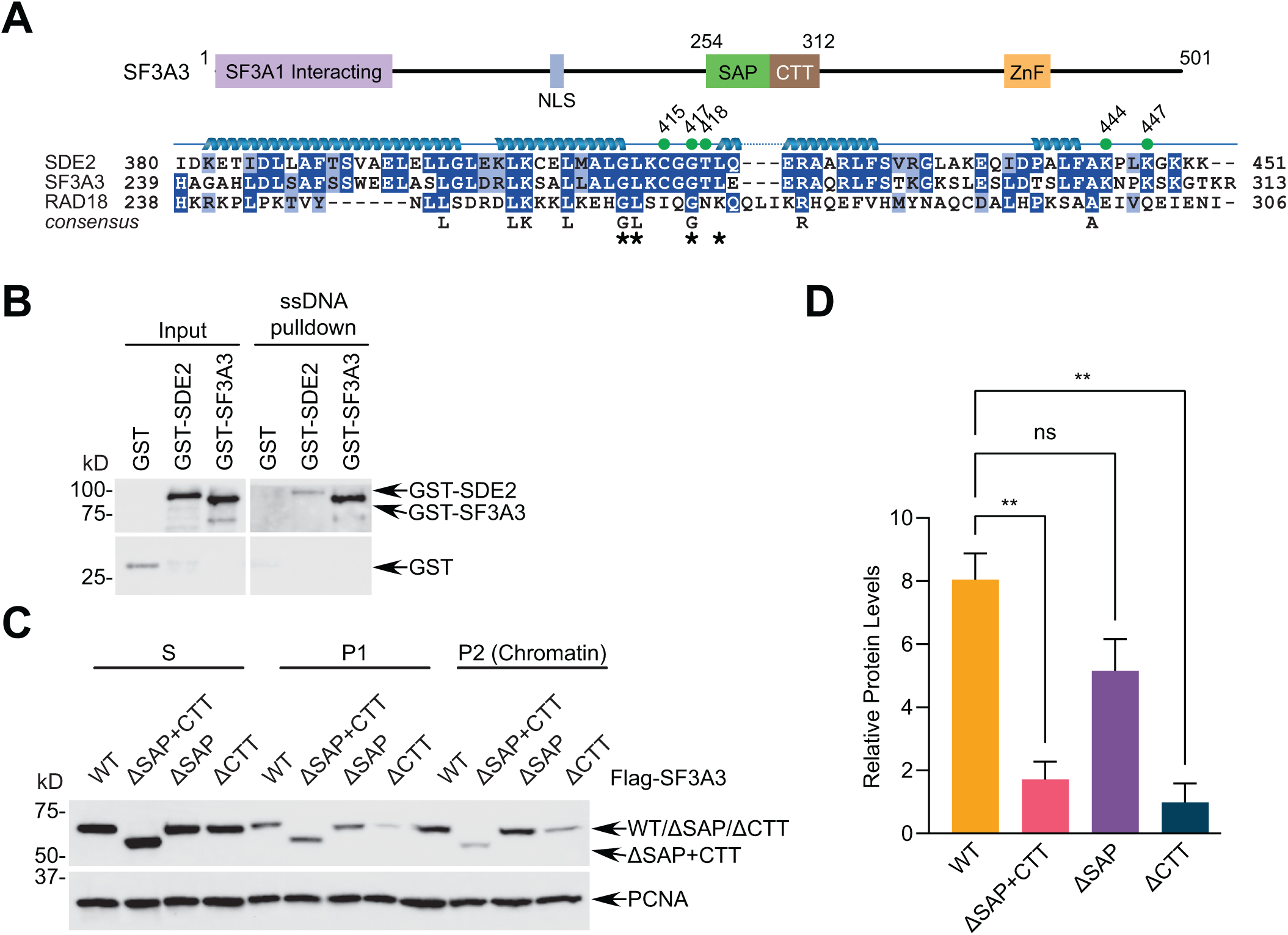
SDE2 and SF3A3 share the unique, extended SAP+CTT motif with similar DNA binding properties. **(A)** Schematic showing the domain structure of SF3A3 and an alignment of the SAP+CTT region of SDE2, SF3A3, and RAD18. Green dots indicate residues identified from ^1^H-^15^N HSQC, and asterisks indicate the residues mutated in the RAD18-based point mutant (SDE2 GLGL) **(B)** Purified GST-SDE2-Flag WT or GST-SF3A3 was incubated with a biotinylated 80-mer ssDNA oligo for DNA pull-down and analyzed by Western blotting. **(C)** U2OS cells were transfected with Flag-SF3A3 WT, ΔSAP+CTT, ΔSAP, or ΔCTT, fractionated to separate cytosolic and chromatin-associated protein pools, and analyzed by Western blotting. **(D)** Quantification of three independent experiments; error=SEM, \*\**p*<0.01, two-way ANOVA.

### SDE2^SAP^ is required for supporting TIM-dependent replication fork progression

Lastly, we determined how the ability of SDE2 to interact with DNA via its SAP domain is linked to its role in DNA replication. We previously showed that the SDE2 domain (SDE2^SDE2^), the conserved coiled-coiled domain of SDE2 at its N-terminus, directly interacts with TIM to support the localization and stability of TIM at replication forks, thus ensuring the functional integrity of the FPC in the replisome (19). We reasoned that the ssDNA-specific interaction of SDE2^SAP^ may help tether TIM at replication forks, further stabilizing the replisome required for fork progression. Here, we employed the Retro-X Tet-On system, where we could reconstitute SDE2 knocked-down cells with siRNA-resistant SDE2 WT or ΔSAP mutant in a doxycycline (dox)-inducible manner (**Fig. 7A**). Subcellular fractionation revealed that the localization of the ΔSAP mutant at the P2 chromatin fraction is dramatically impaired with a concomitant increase of its presence in S and P1 fractions, compared to wild-type (**Fig. 7B**). Importantly, a TIM-EdU PLA assay to visualize TIM at EdU-labeled ongoing replication forks demonstrated that unlike the wild-type, the ΔSAP mutant fails to complement the percentage of cells with positive PLA foci to the level of control following siRNA knockdown and dox induction, indicating that SDE2^SAP^ is necessary for the localization of TIM at replication forks, similar to the phenotype of the ΔSDE2 domain mutant as we previously showed (**Fig. 7, C and D**) (19). Accordingly, DNA combing analysis, which allows for monitoring dynamics of single DNA replication tracks, revealed that cells re-expressing the ΔSAP mutant exhibit a significant shortening of replication tracks in comparison to wild-type cells, indicating that SDE2^SAP^ is required for the function of TIM to support efficient fork progression (**Fig. 7E**). Similar defects in TIM localization and fork progression were observed in the SAP CGTKK point mutant (**Fig. S7**, **A-C).** Together, these results suggest that the DNA binding property of SDE2 mediated by SDE2^SAP^ is essential for its function at DNA replication forks, ensuring the integrity of the FPC and fork stability via TIM regulation.

**Figure 7.**
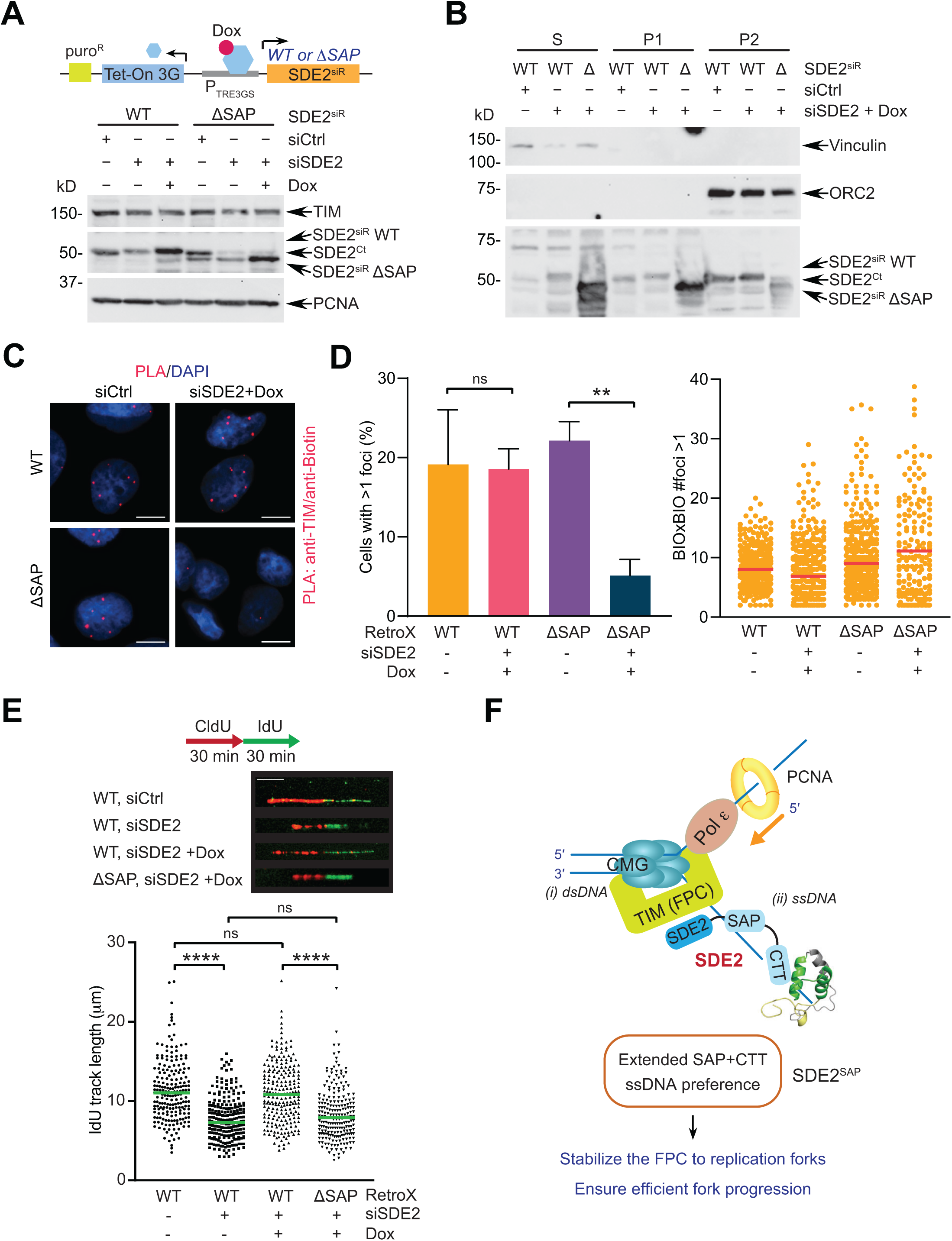
The ssDNA binding ability of SDE2 is required for the function of the fork protection complex (FPC) at replication forks. **(A)** Top: Retro-X Tet-One system. Doxycycline (dox) induces the expression of siRNA-resistant cDNA under the P_TRE3GS_ promoter upon its binding to the Tet-On 3G transactivator. Bottom: induction of SDE2 WT or ΔSAP (Δ395-451) in response to dox, following siRNA transfection. **(B)** Subcellular fractionation of U2OS cells re-expressing SDE2 WT or ΔSAP into S (cytosolic), P1 (nuclear, non-chromatin), and P2 (chromatin) fractions. **(C)** Representative images of TIM:EdU PLA foci in the Retro-X SDE2 WT or ΔSAP cells following SDE2 siRNA transfection and dox induction. Scale bar: 10 μm. **(D)** Left: quantification of cells positive for TIM-EdU PLA foci (>400 cells per condition, n=3, error=SEM, \*\**p*<0.01, t-test). Right: number of biotin-biotin foci (controls for marking active replication forks) per PLA-positive cell; line=median. **(E)** Top: Representative images of DNA fibers labeled with CldU and IdU from the Retro-X SDE2 WT or ΔSAP cells. Bottom: dot plots of the DNA fiber IdU track length (>200 tracks per condition, n=2, \*\*\*\**p*<0.0001, ns, not significant, Mann-Whitney). **(F)** A model depicting the role of SDE2^SAP^ in DNA replication via TIM regulation. We propose that together with the FPC gripping dsDNA in the front of the replisome (i), ssDNA-specific binding of the extended SAP domain of SDE2 contributes to tethering of TIM at ongoing replication forks (ii), further stabilizing replisome association and promoting efficient fork progression.

## DISCUSSION

### Roles of SDE2^SAP^ in DNA replication fork integrity

In this study, we present structural, biochemical, and biological evidence that the SDE2 SAP domain, SDE2^SAP^, is a bona fide ssDNA binding motif that is essential for the function of SDE2 at DNA replication forks. As a PCNA-associated regulatory component of the replisome, SDE2 interacts with TIM in the FPC and stabilizes TIM against proteolytic degradation (19). The conserved SDE2 domain, SDE2^SDE2^, directly mediates its binding to TIM, and disruption of the SDE2-TIM interaction impairs proper localization of TIM to sites of DNA replication, causing disruption of replication fork progression and loss of stalled fork protection under replication stress. We propose that, besides SDE2^SDE2^, SDE2^SAP^ constitutes an essential element for TIM regulation by tethering SDE2 in the FPC to DNA replication forks (**Fig. 7F**). Consequently, mutations in SDE2^SAP^ and subsequent loss of its DNA binding capacity decrease the affinity of TIM to replication forks and impede DNA replication fork progression. Intriguingly, previous structural studies revealed preferential interactions of the Tof1-Csm3 complex to dsDNA over ssDNA by gripping dsDNA at the front of the replisome to stabilize it (12, 33). Hence, it is plausible that while the FPC is engaged with dsDNA, SDE2^SAP^ is specialized for increasing the affinity of the FPC to ssDNA-exposed replication forks. TIPIN, the heterodimeric partner of TIM, is known to stabilize the FPC on the stretch of RPA-coated ssDNA at stalled forks and promote damage-inducible CHK1 phosphorylation via its direct interaction with RPA, and SDE2 may play a primary role in ssDNA association when RPA accumulation is minimal in unchallenged conditions (16). Together, our study highlights the vital role of SDE2^SAP^ in preserving the structural integrity of the FPC, thus promoting replisome progression. It would be interesting to determine whether the affinity of SDE2 to stalled forks changes in response to DNA replication stress, thus modulating the dynamics of the FPC and the replisome as necessary for the protection and recovery of damaged forks.

### Structural diversity of the SAP domain

The canonical SAP domain consists of two amphipathic helices connected by an extended loop region with a series of positively charged and hydrophobic amino acids, including a highly conserved glycine residue in the loop shortly before the second helix. This glycine was one of the residues strongly perturbed in our NMR ^1^H-^15^N HSQC analysis, supporting the idea that the extended loop-helix junction is a major interface for DNA binding, which is consistent with previous reports from Ku70 and RAD18 where an exposed face of the helix bundles constitutes the binding site for DNA (27, 28). While the common DNA binding property was proven by several seminal biochemical and structural analyses of multiple SAPs, each motif showed distinct variations as well. The SAF-A SAP was shown to bind the minor groove of the A-tracts in SARs (scaffold attachment regions) via a mass binding mode, suggesting that SAF-A^SAP^ may exhibit some levels of sequence specificity (34). The SAP domain of Ku70, which exhibits well-defined three helices with a basic N-terminal flexible loop (28), is connected to the C-terminus of the Ku70/Ku80 heterodimeric complex via a disordered linker. Its relatively low DNA binding affinity indicates that Ku70^SAP^ is likely to promote stable association of Ku70/80 to DNA ends and regulate their inward movement at the junction (29,35,36). The SAP domain of PIAS1, an E3-type SUMO ligase, forms a unique four-helix bundle at its N-terminus, a part of which constitutes a helix-extended loop-helix SAP motif (i.e. α2- and α3-helices) (37). PIAS1^SAP^ was also shown to bind A/T-rich DNA oligomers, sharing a similar function of scaffold attachment proteins in active transcription regions. These findings suggest that while the core helix bundle is a major DNA recognition motif of SAP, variable structure- or sequence-specific DNA binding modes may exist among distinct SAP domains.

Indeed, the NMR structure we resolved in this study reveals an extended SAP feature that harbors a unique C-terminal tail following the core SAP fold, designated as SDE2^CTT^, highlighting the structural diversity within the SAP domain family. Loss of both SDE2^SAP^ and SDE2^CTT^ is required for complete abrogation of SDE2 binding to DNA, indicating that SDE2^CTT^ augments the ssDNA binding mediated by a canonical SAP domain. In particular, two lysine residues identified from the HSQC analysis are expected to stabilize the interactions with the phosphate backbone of DNA. Intriguingly, the extended configuration of SDE2^SAP+CTT^ is also observed in the middle region of the human splicing factor subunit SF3A3. The sequence similarities of SAP domains are the highest between SDE2 and SF3A3, versus SDE2 compared to other SAP domains such as RAD18, Ku70, or PIAS. This high similarity includes C415, G417, T418, K444, and K447, the key residues revealed from the HSQC analysis, indicating that these two domains may have been evolutionarily selected as a functional unit in both proteins.

### The ssDNA binding mode of SDE2^SAP^

The specificity of SDE2^SAP^ for ssDNA implies that it may be optimized for association with replication forks. Over five-fold higher affinity to ssDNA versus dsDNA was previously reported in the SAP domain of RAD18, which is known to associate with RPA-coated ssDNA (27). Our results indicate that SDE2^SAP^ and RAD18^SAP^ share many properties, including similar ssDNA binding affinity (∼1.0 μM) and interacting interfaces, further supporting the specialized role of SAP in regulating the replication stress response.

There are several types of ssDNA binding motifs present in DNA repair proteins. RPA, a heterotrimer of RPA70, RPA32, and RPA14, possesses six OB (oligonucleotide/oligosaccharide binding) fold domains, four of which act as ssDNA binding domains (38). RPA is known to occupy ∼30 nucleotides per trimer with high affinity (∼10^-4^ μM) (39). Structural analyses revealed that flexible structural loops in each OB-fold, which keeps a binding pocket open, clamp down on DNA and stabilize the interaction in the closed conformation upon ssDNA binding, in which basic amino acids make hydrogen bonds with the DNA phosphate while aromatic side chains stack with DNA bases (40, 41). Conserved basic residues in SDE2^SAP^ may exert a similar role during DNA contact. The HIRAN domain present in HLTF, a SWI/SNF family DNA translocase involved in stalled fork reversal (42), constitutes a modified OB fold that specifically recognizes 3’ ssDNA ends (43). The free 3’-hydroxyl group is nestled deep in the back of the pocket, while other hydrogen bonds preclude binding of dsDNA in the pocket and ssDNA is stabilized by electrostatic interactions with arginine and lysine residues; this unique feature of 3’ ssDNA end recognition catalyzes the regression of stalled replication forks. SPRTN, a metalloprotease responsible for resolving DNA-protein cross-links (44), contains a Zn^2+^- binding sub-domain (ZBD) in the catalytic SprT domain, which preferentially binds to ssDNA through an aromatic pocket lined by tyrosine and tryptophan, which is only wide enough to accommodate unpaired bases in ssDNA (45). For SDE2^SAP^, based on the available structures and mutagenesis studies, we propose a cleft formed by both hydrophobic and basic residues at the loop-helix 3 junction and positively-charged residues in the CTT, which together is able to accommodate ssDNA, although a distinct structure of the SAP complex with DNA needs to be experimentally verified in the future. Without any clamp or pocket, the core SAP in general is expected to be relatively low affinity, and the CTT, uniquely present in SDE2, may help stabilize the interaction. This feature has not been reported in any other SAP domains, and closer scrutiny to the surrounding region of other SAP domains may lead to the identification of similar supporting roles.

### The versatile function of SDE2 in DNA and RNA transactions

It is worthwhile to note that SAP domains are also frequently associated with elements involved in the assembly of RNA-processing complexes, and emerging evidence supports that SDE2 may play a role in mRNA splicing and ribosome biogenesis (46). Nevertheless, whether the SAP domain is directly involved in binding RNA remains elusive. The *S. pombe* Sde2, which lacks the SAP domain, is known to regulate pre-mRNA splicing by promoting the association of Cactin into spliceosomes (25). The cryo-EM structure of the human post-catalytic spliceosome also revealed that the N-terminal region of SDE2, not the C-terminal SAP, stabilizes Cactin into the spliceosome via its interaction with CRNKL1 (47). Notably, SDE2 is shown to associate with non-coding RNAs, though whether SDE2^SAP^ is required for this interaction remains unexplored (46). In our hands, recombinant SDE2 fails to form a protein-RNA complex *in vitro*, although we do not exclude the possibility that SDE2 recognizes specific structures of rRNA or snoRNA in association with the riboprotein complex (**Fig. S7D**). The CTTs from both human SDE2 and SF3A3, located adjacent to the SAP motif, are likely a defunct RNA binding zinc finger from yeast (**Fig. S6**), raising the possibility that both may have evolved to acquire the ability to bind DNA. Gain of the SAP domain in metazoans may have diversified the function of SDE2, allowing its roles in DNA replication and the DNA damage response alongside its primordial role in RNA transactions.

In conclusion, our study reveals the existence of a previously uncharacterized extended SDE2^SAP+CTT^ domain and elucidates its role in guiding SDE2 function at replication forks via its ssDNA binding property. A high degree of SAP conservation throughout evolution and its enrichment in genome maintenance factors may provide important clues for the discovery of new DNA repair and damage response factors and for their functional studies.

## EXPERIMENTAL PROCEDURES

### Cell culture and plasmid construction

U2OS and HEK293T cells were acquired from the American Tissue Culture Collection (ATCC). Cells were grown with Dulbecco’s Modified Eagle Medium supplemented with 10% fetal bovine serum and 1% v/v Penicillin/Streptomycin, and incubated at 37 °C in a humified chamber under 5% CO_2_. pcDNA3-SDE2-Flag and its mutants were previously described. Full-length SDE2 was subcloned into pEGFP-N1 (Clontech) for live-cell imaging, and full-length SDE2 or SDE2 aa380- 451 was subcloned into pGEX-6P1 (GE Healthcare) for recombinant protein production. SDE2 aa78-451 (ΔUBL) was subcloned to pcDNA3-N-Flag to generate N-end rule-refractory SDE2 proteins. For RetroX stable cell line generation, the SDE2 cDNA was subcloned into the retroviral pRetroX-TetOne-puro vector (Clontech). The SF3A3 cDNA was obtained from the DF/HCC DNA Resource Core (PlasmID: HsCD00043137) and cloned into pcDNA3-N-Flag. Primers containing restriction sites were used to amplify cDNAs for subcloning and primers with mutations or deletions were used for site-directed mutagenesis (SDM). Mutations were verified with Sanger DNA sequencing (Stony Brook University Genomic Facility). SDM primer information can be found in **Table S1**.

### DNA and siRNA transfection

DNA transfections were performed with GeneJuice Transfection Reagent (MilliporeSigma), while siRNA-mediated knockdowns were achieved by Lipofectamine RNAiMAX (Thermo Fisher). Target sequences for siRNA-mediated knockdown were 5’-ACGGCAATGGCCTACTAAA-3’ (siSDE2#1) and 5’- GTAGCTTAGTCCTTTCAAA-3’ (siTIM#1), and the siRNA oligonucleotides were synthesized by Qiagen. The AllStars negative control siRNA (Qiagen #SI03650318; CAGGGTATCGACGATTACAAA) was used for control transfection.

### Cell lysis, fractionation, and Western blotting

Cells were harvested with trypsin or PBS, washed, resuspended in NETN300 buffer (50 mM Tris-HCl pH 7.5, 300 mM NaCl, 0.3 mM EDTA, 1% NP-40) complemented with ETDA-free protease inhibitor (PI) cocktail (Roche), incubated on ice for 40 min, centrifuged at 14k rpm, 4 °C for 10 min, and the supernatant saved as whole cell lysate (WCE). For subcellular fractionation, cells were incubated on ice for 20 min, and centrifuged at 14k rpm, 4 °C for 10 min. The supernatant, containing the cytosolic proteins, was saved as the S fraction. The pellet (nuclei) was washed with S buffer, resuspended in P1 low salt buffer (10 mM Tris-HCl pH 7.5, 0.2 mM MgCl_2_, 1% Triton X-100) with PI, incubated on ice for 15 min, centrifuged, and the supernatant, containing nuclear, non-chromatin proteins, was saved as the P1 fraction. The pellet (chromatin) was washed with P1 buffer, resuspended in 0.2 N HCl, incubated on ice for 20 min, centrifuged, and the supernatant transferred into a tube containing an equal volume of Tris-HCl pH 8.0 to neutralize the acid. These acid-soluble, chromatin-associated proteins were saved as the P2 fraction. Protein lysates were resuspended in 2X Laemmli Buffer, boiled, loaded into SDS-PAGE gels, and transferred to PVDF membranes (MilliporeSigma). Membranes were blocked in 5% milk in TBS with 0.1% Tween-20 (TBS-T) for 1 h, then incubated in the indicated primary antibodies in 1% milk overnight at 4 °C. The membrane was washed three times with TBS-T, incubated in HRP-conjugated secondary antibodies at RT for 1 h, then washed three times with TBS-T. HRP signals were detected by enhanced chemiluminescence (ECL) Western blotting substrates (Thermo Fisher) using either HyBlot CL autoradiography film (Thomas Scientific) or using an iBright digital imager (CL1000; Thermo Fisher).

### Antibodies, reagents, and chemicals

Primary antibodies used were anti-Flag (MilliporeSigma F1804, 1:8000), anti-GFP (Santa Cruz SC-9996, 1:1000), anti-GST (GenScript A00865, 1:2000), anti-ORC2 (BD Biosciences 551178, 1:1000), anti-PCNA (PC10; Santa Cruz SC-56; 1:4000), anti-SDE2 (Sigma Atlas HPA031255, 1:4000), anti-TIM (Bethyl A300-961A, 1:2000), and anti-vinculin (H-300; Santa Cruz SC-5573, 1:1000). Secondary antibodies used were anti-mouse IgG HRP (Cell Signaling Technology #7076, 1:4000) and anti-rabbit IgG (Cell Signaling Technology #7074, 1:4000). Chemical and reagent information can be found in **Table S2**.

### Protein expression

GST or GST-SDE2 was expressed using BL21 (DE3) cells by incubating at 37 °C to an OD_600_ of 0.6-0.8, then inducing with 0.5 mM isopropyl β-D-1-thiogalactopyranoside (IPTG, MilliporeSigma) at 30 °C for 6 h. To increase protein solubility, cells were sometimes grown and induced at 16 °C for 15-18 h. After centrifugation, cells were resuspended in PBS with 1 mg/mL hen egg white lysozyme and 0.5 mM phenylmethylsulfonyl fluoride (PMSF), rocked at 4 °C for 40 min, and stored at -80 °C. After thawing on ice, the mixture was sonicated on a QSonica Q500 Digital Sonicator at 50% amplitude with three cycles of 10 s pulses followed by 20 s recovery time.

Triton X-100 was added to a final concentration of 0.5-1%, and the lysate was rocked for 30 min at 4 °C. Lysates were centrifuged at 14k rpm for 15 min at 4 °C, then filtered through a 0.45 μm PES filter. Cleared lysates were either purified immediately or frozen at -80 °C in aliquots. Recombinant GST-SF3A3 was obtained from Novus Biologicals (H00010946-P01).

### Protein purification

Recombinant proteins were purified using the batch method, gravity columns, or an FPLC. For batch purification, 250-1000 μL cleared lysate was thawed on ice and added to 50-100 μL of prepared Glutathione Agarose Affinity resin (Thermo Fisher), then incubated at 4 °C while rocking for 2-4 h. The resin was washed three times with wash buffer (50 mM Tris-HCl pH 8.0, 150 mM NaCl, 0.1 mM EDTA). If the GST tag was to be removed, the resin was washed once with PBS and twice with protease buffer (50 mM Tris-HCl pH 8.0, 150 NaCl, 1 mM DTT, 1 mM EDTA), then 8 units of HRV 3C protease (Thermo Fisher) was added per 50 μL resin and rocked at 4 °C overnight. The supernatant containing the cleaved protein was then recovered. Otherwise, the protein was eluted with 50 mM Tris-HCl pH 8.0 containing 10 mM reduced glutathione (GSH) while rocking at 4 °C for 30 min. Elution was repeated once and both eluates combined. If necessary, protein was dialyzed 2000-fold overnight with dialysis buffer (50 mM Tris-HCl pH 8.0, 1 mM DTT, 0.2 mM EDTA, 10% v/v glycerol). For gravity purification, 1 mL of resin was packed into a Poly-Prep Chromatography Column (Bio-Rad) and washed with 5 column volumes (CV) PBS, 5 CV PBS-T, 10 CV wash buffer. 500-1000 μL of lysates were loaded into the closed column. Wash buffer was added to bring the final volume loaded to 5-9 mL and the column was capped and rocked overnight at 4 °C. The next morning the resin was allowed to settle and the flow-through (FT) was collected. The resin was washed twice with 5 CV of wash buffer while rocking at 4 °C for 15 min. Finally, the protein was eluted in 0.5 mL fractions over 3 CV with elution buffer (wash buffer with 10 mM GSH). Fractions containing the protein-of-interest (POI) were pooled, and either dialyzed overnight with dialysis buffer or concentrated with Amicon centrifugal filters (MilliporeSigma) with appropriate molecular weight cut-offs. For FPLC purification, a 5 mL SP FF column (cation exchange column; GE Healthcare) was attached to an AKTA chromatography system and equilibrated with 5 CV buffer SA (20 mM Tris-HCl pH 7.5, 50 mM NaCl, 1 mM DTT or TCEP, 5% v/v glycerol), 5 CV buffer SB (buffer SA + 1 M NaCl), and 5 CV buffer SA, then cleared lysate with diluted to ∼40-45 mL in buffer SA, 0.5 mM PMSF was added, and the sample was loaded onto the column at 3 mL/min while recycling the FT until all protein was loaded as determined by the UV trace. The column was washed with 5 CV buffer SA, and then the protein was eluted over a linear salt gradient of 0-100% buffer SB over 10 CV at 1-3 mL/min. The fractions were run on an SDS-PAGE, and appropriate fractions were pooled and loaded onto a 5 mL GSTrap FF column (affinity column; GE Healthcare) that had been equilibrated with 5 CV PBS and 5 CV buffer SA+100 (buffer SA + 100 mM NaCl) at 0.1-0.5 mL/min while recycling the FT. The column was washed with 5 CV buffer SA+100, and the protein was eluted with elution buffer (buffer SA+100 + 10 mM GSH) in 0.5 mL fractions at 0.1-0.5 mL/min over 3 CV. Fractions containing the POI were pooled and 10% v/v glycerol was added.

### Electrophoretic mobility shift assay

To create the desired DNA structures, a 5’ 6-carboxy fluorescein (FAM)-labeled oligo and an unlabeled complement were combined at equimolar concentrations in annealing buffer (10 mM Tris-HCl pH 7.5, 50 mM NaCl, 1 mM EDTA), heated to 95 °C in a heat block, allowed to cool to room temperature (RT), and stored at -20 °C in amber-colored tubes. For the binding reactions, 10-50 nM of DNA was incubated with increasing concentrations of purified, recombinant protein in Tris binding buffer (10 mM Tris-HCl pH 7.5, 50 mM NaCl, 0.5 mM DTT, 0.05 mM EDTA, 3% v/v glycerol, 0.05 mg/mL BSA) or phosphate binding buffer (50 mM Na/K PO_4_ pH 6.0, 50 mM NaCl, 5 mM DTT) on ice for 10-30 min. While incubating the binding reaction, a 3-6% TBE-based, native PAGE gel was pre-run at 4 °C, 200 V in 0.5X TBE. Before loading the samples, either orange G, xylene cyanol FF, or an orange G/xylene cyanol FF hybrid loading dye was added, and the samples were run at 170-200 V for 30-60 min. The gel was left in the glass plates, was dried and cleaned, and directly imaged using a Typhoon FLA 9000 phosphorimager set to fluorescence for FAM with a PMT of 500, utilizing the multistage. Images were processed and quantified using Fiji. For the supershift assays, the following antibodies were added to the reactions and incubated for an additional 15 min before loading into the gel: anti-Flag (MilliporeSigma F1804), anti-SDE2 (Sigma Atlas HPA031255), or anti-αtubulin (B-7; Santa Cruz SC-5286). FAM-labeled and unlabeled oligonucleotide sequences can be found in **Table S3**.

### Chemical shift perturbation

SDE2^SAP+CTT^ was purified in-house using the above procedure, but grown with M9 minimal media; 1X M9 salts [47.75 mM Na_2_HPO_4_·7H_2_O, 22.04 mM KH_2_PO_4_, 8.56 mM NaCl], 2 mM MgSO_4_, 18.35 mM ^15^NH_4_Cl, 1X Solution Q [10 mg/L FeCl_2_·4H_2_O, 368 ng/L CaCl_2_·2H_2_O, 128 ng/L H_3_BO_3_, 36 ng/L CoCl_2_·6H_2_O, 8 ng/L CuCl_2_·2H_2_O, 680 ng/L ZnCl_2_, 12.1 ng/L Na_2_MoO_4_·2H_2_O, 96.8 mN HCl], 22 mM D-glucose [^12^C for single-labeled samples, ^13^C for double-labeled samples], 1X vitamin mix [5 mg/L thiamine-HCl, 1 mg/L D-biotin, 1 mg/L cholic acid, 1 mg/L folic acid, 1 mg/L niacinamide, 1 mg/L D-pantothenate, 1 mg/L pyridoxal 5’-phosphate, 100 ng/L riboflavin], 100 mM CaCl_2_, and 1X ampicillin. Samples were buffer exchanged into 50 mM Na/K phosphate, 50 mM NaCl, 5 mM DTT, 10% D_2_O; the final concentration of the protein was 200 μM. To show chemical shift perturbations upon binding to ssDNA, ^1^H-^15^N HSQC-TROSY spectra were gathered using the standard Bruker pulse sequence, trosyf3gpphsi19.2 (NS: 16; TD: F1-1H 1024/ F2-15N 128; SW: F1-1H 12.00/ F2-15N 36.00). In all cases, the chemical shift perturbations values (Δδ, in ppm) were calculated using Equation:

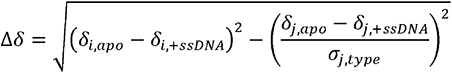

where *i* and *j* index the ^1^H (amide) and the corresponding heteronucleus (^15^N or ^13^C), respectively. Spectra were gathered for SDE2^SAP+CTT^ apo form, then a 16-mer oligo of random sequence but designed to prevent self-annealing and hair-pinning (GCTATGGAGAACGGTA) was added to the sample at a 1:1 protein to ssDNA ratio and the same spectra were gathered; the process was repeated for ratios of 1:2 and 1:4. The four spectra were then overlaid to identify chemical shift perturbation induced by ssDNA.

### NMR structure determination

For the structural calculation of SDE2^SAP+CTT^, spectra were acquired using the standard Bruker pulse sequence. Detailed spectra information can be found in **Table S4**. All spectra were acquired with 200 μM SDE2 without ssDNA. All ^1^H-^15^N-^13^C (except CCCONH) and ^1^H-^15^N heteronuclear NMR experiments were acquired at 25 °C on a Bruker Avance III HD spectrometer operating at a ^1^H frequency of 700 MHz equipped with an Inverse Triple Resonance (TXI) 5 mm CryoProbe. With the same sample, CCCONH, NOESY, and IPAP experiments were acquired at 25 °C using a Bruker Avance III HD spectrometer operating at a ^1^H frequency of 850 MHz equipped with a Triple Resonance (TCI) ^13^C-enhanced 5 mm CryoProbe. Sample stability was investigated by ^1^H NMR experiments (zgesgp; NS:8) prior and post all 2D/3D NMR experiments. The spectra were processed using Bruker TopSpin 3.6.3 and backbone assignments were analyzed and established using NMRFam Sparky and its plug-in webserver I-PINE (48); these assignments were confirmed using NMRViewJ (49). Structure calculations for SDE2 was performed using the ARIA2.3 suite (50, 51), utilizing distance restraints, experimentally obtained from ^13^C-edited NOESY-HSQC and ^15^N-edited NOESY-HSQC, supplemented with backbone dihedral angle restraints obtained from the chemical shift statistics using the TALOS-N suite (52). The PARALLHDG force field with PROLSQ for non-bonded parameters was used (53). A simulated annealing protocol comprising of 20,000 steps (27 fs integration time) was carried out at high temperature (10,000 K) followed by two Cartesian cooling phases of 1,000 K and 50 K of 40,000 steps each (3 fs integration times). A network anchoring protocol was introduced for the first three iterations of the protocol using default parameters, and floating chirality was implemented for prochiral moieties. Starting with the fourth iteration, hydrogen bonding restraints were added for residues predicted by TALOS-N to be part of defined secondary structure elements. The final production run consisted to a similar simulated annealing protocol as described above except that 1000 structures were generated for each iteration. At the final step, the 100 lowest energies structures were submitted to a short restrained molecular dynamics simulation in explicit solvent using XPLOR-NIH (54). For the final NMR ensembles, the 20 lowest energies structures displaying the lowest RDC Q values (see below) that showed no distance restraint violations larger than 0.5 Å and no dihedral angle violations larger than 5° were analyzed using PROCHECK-NMR (55), wwPDB and MOLPROBITY (56). The final structural ensembles, comprising 20 structures was deposited in the PDB with accession codes 7N99.

### Residual dipolar couplings and backbone dynamics

Residual dipolar couplings (RDCs) were measured utilizing aligned media generated by the direct addition of Pf1 phage (ASLA Biotech) into the NMR samples of SDE2 (400 µM) to final phage concentrations of 14.7 mg/mL. In order to optimize the degree of alignment for the NMR sample, NaCl concentrations was adjusted to reach a final concentration of 108 mM. Amide ^15^N,^1^H RDC values were extracted from a set of 2D HSQC-IPAP (57) experiments (512 and 300 complex points using sweep-widths of 12 ppm and 46 ppm in the ^1^H and ^15^N dimensions, respectively). The alignment tensor values were obtained using PALES (58). Only regions with well-defined secondary structure values were selected. The Q values were defined as follow (59):

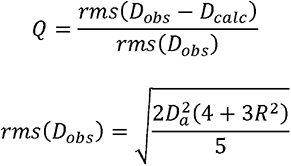

Where D_obs_ and D_calc_ are the observed and calculated values of the ^15^N,^1^H RDC values; Da and R are the anisotropy and rhombicity of the alignment tensor.

A backbone dynamics experiment, steady-state NOESY (ssNOE), was used to measure the backbone motion of each residue. The TALOS-predicted dihedral angles were selected based on the ssNOE ratio using 0.7 as the cutoff, while the TALOS-predicted dihedral angles of the residues with a ssNOE ratio<0.7 were discarded as the backbones were too flexible. The TALOS angle restraints and RDC restraints were used to validate initial structure calculation.

### Biotin-DNA pulldowns

To create the DNA, a 5’ biotinylated 80-mer was mock annealed or annealed with an unlabeled complement at a 1:4 molar ratio in annealing buffer (10 mM Tris-HCl pH 7.5, 50 mM NaCl, 1 mM EDTA) by heating to 95 °C in a heat block, allowed to cool to RT, and stored at -20 °C. Pierce streptavidin magnetic beads (Thermo Fisher) were prepared by washing twice with TBS- T, and equilibrated with oligo-binding buffer (20 mM Tris-HCl pH 7.5, 500 mM NaCl, 1 mM EDTA). For each reaction, 5 μL of beads was incubated with 10 pmol of DNA in oligo-binding buffer at RT for 30 min while rocking, then washed with oligo-binding buffer three times. Purified protein was added to the beads in protein-binding buffer (10 mM Tris-HCl pH 7.5, 100 mM NaCl, 0.2 mM EDTA, 1 mM DTT, 10% v/v glycerol, 0.01% NP-40, 10 μg/mL BSA), rocked for 20-30 min at RT, washed with protein-wash buffer (protein-binding buffer with 100 mM NaCl) four times while rocking for 3 min each. Beads were boiled in 20 μL Laemmli buffer for 15 min to break the crosslinker, and the supernatant was run in an SDS-PAGE gel and analyzed by Western blotting.

### Fluorescence anisotropy

GST-tagged recombinant full-length SDE2 or SDE2^SAP+CTT^ and FAM-labeled oligo were combined in 96-well, black fluorimetry plates with clear bottoms (Greiner Bio-One) to a final concentration of 1X buffer (50 mM Na/K PO_4_, 50 mM NaCl, 5 mM DTT, pH 6.0), 1 μM DNA, and varying, known concentrations of protein from 0-32 μM. Each protein concentration was set up in duplicate. The plate was read using a SpectraMax M5 Microplate Reader (Molecular Devices) in polarization mode, with the photomultiplier tube set to ’medium’ and 100 flashes per read. The excitation and emission wavelengths were set to 492 nm and 518 nm, respectively, with a cutoff filter at 515 nm. The raw data was reduced using the standard anisotropy calculation and a G factor of 1. Anisotropy values were plotted against protein concentration and fit to a total binding curve, single site in Prism 8/9 (GraphPad).

### Fluorescence Loss in Photobleaching (FLIP)

Cells were seeded 24-48 h before imaging in 35 mm dishes with a 14 mm microwell using No. 1.5 coverglass (MatTek P35GC-1.5-14-C poly-d-lysine coated) at a density of 300,000 and reverse transfected with 1 μg of plasmid DNA using GeneJuice (MilliporeSigma). All constructs had an eGFP tag. Experiments were performed on a Carl Zeiss LSM 510 Meta confocal microscope with a live-cell imaging setup (37 °C, 5% CO_2_) using a 488 nm laser and ×63 objective. Scan area was set to 512x512 px. For each imaged cell, a 47x47 px ROI was defined and a time series was used to capture the rate of photobleaching. After acquiring three pre-bleach images, the laser was illuminated at 10% transmission for 10 iterations to bleach the ROI, repeating for 40 cycles or until the nucleus was completely bleached, imaging after every bleaching event. Time lapse between each image was 3.39 s. Fluorescence intensities at each time point were quantified using Fiji or Zeiss LSM Image Examiner, and these intensities were normalized as a function of the average pre-bleach intensity, with the pre-bleach value defined as 100% and 0 defined as 0%. The normalized values were fit to a two-phase decay model and plotted against time. The curve fits were compared using a two-tailed unpaired t test. All analysis was done using Prism (GraphPad).

### Generation of Retro-X Tet-On inducible cell lines

The retroviral plasmid pRetroX-TetOne puro was acquired from Clontech and amplified using NEB Stable competent *E. coli* (high efficiency). siRNA-1 resistant SDE2 WT or ΔSAP (deletion of amino acids 395-451 along with ΔUBL 1-77) were subcloned into pRetroX-TetOne puro vector. Retroviruses were produced using the GP2-293 packaging cell line (Clontech), where pRetroX-TetOne puro empty vector (EV), SDE2 WT or ΔSAP constructs were co-transfected with the envelope vector pCMV-VSV-G, using Xfect^TM^ transfection reagent (Clontech). U2OS cells were transduced for 16 h using 8 μg/mL polybrene (Sigma-Aldrich). Puromycin selection (2 μg/mL) began 48 h post-infection and lasted for 3 days, until all non-transfected cells had died. After a week, cells had recovered and were tested to find the optimal doxycycline concentration to induce the transgene.

### In situ Protein Interaction with Nascent DNA Replication Forks (SIRF)

For the Flag-SDE2 SIRF, cells were seeded on washed coverslip in a 12-well dish, and reverse transfected with plasmid DNA using Genejuice (MilliporeSigma). Cells were pulsed with 125 μM EdU in pre-equilibrated media for 12 min and stopped with ice cold PBS. For the TIM SIRF, Retro-X cells were reverse transfected with siRNA oligonucleotides using RNAiMAX (ThermoFisher), induced with doxycycline (10-100 ng/mL) at 18 h, refreshed with new doxycycline and seeded on coverslips at 48 h, and pulsed with EdU at 72 h. Coverslips were fixed with 4% paraformaldehyde at RT for 10 min, washed with PBS, and stored in PBS at 4 °C protected from light. On the day of the experiment, the coverslips were washed again, permeabilized with 0.3% Triton X-100 for 3 min at 4 °C, washed three times with PBS, and blocked with 1% BSA at RT for 10 min. Biotin was conjugated to the azide group of EdU using the Click-iT click chemistry kit (Thermo Fisher) according to the manufacturer’s protocol. In brief, Click-iT reaction buffer, 2 mM CuSO_4_, 10 μM biotin-azide, and Click-iT buffer additive were added in order and 30 μL of the mixture was added to each coverslip, which were then incubated in a light-protected, humidified chamber at RT for 30-60 min. Coverslips were washed with PBS, and proximity ligation assay was performed using the DuoLink In Situ PLA & Detection kit (MilliporeSigma) according to the manufacturer’s protocol. All incubations took place in a light-protected, humidified chamber at 37 °C. In brief, coverslips were blocked with 40 μL of blocking solution and incubated for 1 h, incubated in 25 μL of the primary antibodies for 1 h (rabbit anti-TIM, Bethyl A300-961A, 1:500; rabbit anti-Biotin, Bethyl A150-109A, 1:3000; mouse anti-Biotin, Jackson ImmunoResearch 200-002-211, 1:2000), incubated with the PLA plus and minus probes (secondary antibodies) for 1 hr, incubated with the ligase for 30 min, incubated with the rolling polymerase for 1 h 40 min, and mounted using the provided *in situ* wet mounting medium, containing DAPI. The slides were imaged on a Nikon Eclipse Ts2R-FL microscope equipped with a Nikon DSQi2 digital camera and an LED light source, using a ×60 objective. The DAPI channel was visualized with an Ex395/25 Dm425 Em440lp filter set, and the Texas Red channel was visualized using an Ex560/40 Dm585 Em630/75 filter set. For each condition, >400 cells were analyzed. Both imaging and counting were performed with an Eclipse Ts2R-FL inverted fluorescence microscope (Nikon) equipped with a Nikon DSQi2 digital camera and analyzed using the Nikon NIS-Elements BR software and Prism (GraphPad).

### DNA Combing

Exponentially growing cells were pulse-labeled with 50 μM CldU for the indicated time, washed three times with PBS, then pulse-labeled with 250 μM IdU for the indicated time. Cells were harvested by trypsinization, then 400,000 cells pelleted and washed with PBS. DNA fibers were prepared using the FiberPrep® DNA extraction kit and the FiberComb^®^ Molecular Combing System (Genomic Vision, France), following manufacturer’s instructions. In brief, the cells were washed again with PBS before being embedded in low-melting point agarose and cast in a plug mold. After full solidification, plugs were digested overnight with proteinase K. Next day, the plugs were extensively washed prior to short melting and agarose digestion. The obtained DNA fibers were combed onto silanized coverslips (Genomic Vision, France) that were subsequently baked for 2 h at 60 °C. DNA was denatured for 8 min using 0.5 M NaOH in 1 M NaCl. Subsequent immunostaining incubations were performed in humidified conditions at 37 °C. In short, coverslips were blocked with 1% BSA for 30 min, then two primary antibodies were diluted in 1% BSA (rat monoclonal anti-BrdU for CldU, 1:25, and mouse monoclonal anti-BrdU for IdU, 1:5) and incubated for 1 h. After washing the coverslips with PBS-Tween 0.05% (PBS- T), two secondary antibodies were diluted in 1% BSA (Alexa Fluor 594 goat anti-rat and Alexa Fluor 488 goat anti-mouse, 1:100) and incubated for 45 min. Coverslips were washed with PBS- T, dehydrated, and mounted onto microscopic glass slides using ProLong^TM^ Gold Antifade overnight. DNA fibers were imaged with an Eclipse Ts2R-FL inverted fluorescence microscope (Nikon) equipped with a Nikon DSQi2 digital camera and analyzed using Fiji and Prism (GraphPad).

### RNA EMSA

The LightShift Chemiluminescent RNA EMSA kit (ThermoFisher #20158) was used according to the manufacturer’s instructions, in an RNase-free environment. Unless otherwise noted, reagents and materials are included with the kit. In brief, a 6% 0.5X TBE native PAGE gel was pre-run at 100 V in 0.5X TBE running buffer. While pre-running, the binding reactions were prepared using reagents supplied with the kit: either the control protein (iron-responsive protein, IRP) or the experimental protein were incubated with 2-6 μg tRNA, 6.25 nM 3’ biotin-labeled IRE (iron-responsive element) RNA (5’-UCCUGCUUCAACAGUGCUUGGACGGAAC-3’ – biotin, hairpin RNA), and, for the competitive reactions, unlabeled IRE RNA (5’- UCCUGCUUCAACAGUGCUUGGACGGAAC-3’). The 20 μL reactions were incubated at RT for 30 minutes, then mixed with 5X loading buffer and 20 μL loaded into the gel. The gel was run at 100 V until the dye had migrated ½ - ¾ of the way down the gel (∼30-60 minutes). While the gel was running, a nylon membrane (Biodyne 0.45 μm, ThermoFisher #77016) was soaked ≥10 minutes in 0.5X TBE, and the nucleic acid blocking buffer and 4X wash buffer were warmed at 37 °C to dissolve particulate matter. The RNA was transferred to the membrane at 400 mA for 30-45 minutes (until the dye was completely transferred). Excess buffer was removed from the membrane, and the damp membrane was immediately crosslinked at 120 mJ/cm^2^ using a Stratalinker UV Crosslinker 1800 equipped with 254 nm bulbs. Signal was detected using the Chemiluminescent Nucleic Acid Detection Module (ThermoFisher #89880) included with the REMSA kit. In brief, the membrane was blocked with blocking buffer for 15 minutes with gentle orbital shaking, incubated with a 1:300 solution of streptavidin-HRP conjugate in blocking buffer for 15 minutes, rinsed with 1X wash solution, then washed four times with wash solution for five minutes each. The membrane was incubated with substrate equilibration buffer for 5 minutes, blotted to remove excess buffer, then incubated in the chemiluminescent substrate (equal parts luminol/enhancer solution and stable peroxide solution) for 5 minutes without shaking. Excess solution was removed and the membrane was imaged using an iBright 1500 digital imager (CL1000; ThermoFisher) in chemiluminescent mode.

### AlphaFold Prediction

The SDE2 SAP structure (the SAP core + the CTT) was predicted base on primary sequences by AlphaFold2 webserver using the following settings: MSA mode = MMseq2 (Uniref + environmental); model type: auto; paired methods: unpaired + paired; 3 cycles (60).

### Docking Simulation

The model of the extended SDE2 SAP domain that has the lowest energy and most Ramachandran favored regions among the 20 best models was chosen as a receptor protein. The ligand ssDNA used was obtained from the crystal structure of MoSub1-ssDNA complex (PDB ID#: 5ZG9) (61). One hundred simulation models of SDE2 SAP bound to ssDNA were generated by the HDOCK webserver (62, 63) using the template-based docking method. The residues previously identified by NMR (amino acids 415-418, 444, and 447) were defined as receptor binding site residues during the docking simulation. The best docking model was chosen based on the energy and model similarity. The top five models with the lowest energy except models 1 and 4 suggested similar binding behavior between ssDNA and SDE2 SAP, and model 3 was used for figure preparations.

### Sequence Alignments

Reference sequences were obtained from NCBI GenBank (NIH) and trimmed to the desired region if necessary, then aligned using the ClustalΩ algorithm through the Galaxy@Pasteur online computing cluster (Institut Pasteur) with default settings (64). Results were exported as an aligned FASTA file and uploaded to BoxShade (ExPASy/Swiss Institute of Bioinformatics) for shading, then recolored in Adobe Illustrator.

### Statistical Analysis

For all experiments, a minimum of three independent experiments were conducted. Data was analyzed in GraphPad Prism (version 8 or 9), where statistical significance was assessed using either a two-way ANOVA test (chromatin extraction, EMSA, biotin-DNA pulldowns) or a two-tailed Mann-Whitney test (SIRF foci number distribution), with a 95% confidence interval.

## DATA AVAILABILITY

Atomic coordinates and structure factors for the reported NMR structure have been deposited with the Protein Data Bank under 7N99 and the Biological Magnetic Resonance Data Bank under 30927.

*This article contains supporting information*.

## ACKNOWLEDGEMENT

We thank Orlando Schärer (IBS-CGI, Korea) and members of the Kim laboratory for critically reading the manuscript. We thank Francis Picard and Marine Zillow in the Stony Brook University NMR facility for their excellent support. We thank Guowei Tian in the Stony Brook Central Microscopy Imaging Center (CMIC) for training and support in confocal usage for the FLIP studies. We would also like to thank Bruce Johnson (Advanced Science Research Center, CUNY) for verifying the backbone assignments and the calculated SAP structure.

