## Supporting information for "Extended DNA binding interface beyond the canonical SAP domain contributes to SDE2 function at DNA replication forks"

[1]Department of Biochemistry and Cell Biology, State University of New York at Stony Brook, Stony Brook, New York 11794, USA
[2]Department of Chemistry, State University of New York at Stony Brook, Stony Brook, New York 11794, USA
[3]Department of Pharmacological Sciences, State University of New York at Stony Brook, Stony Brook, New York 11794, USA
[4]Department of Chemistry and Biochemistry, The City College of New York, New York 10031, USA
[5]Stony Brook Cancer Center, Renaissance School of Medicine at Stony Brook University, Stony Brook, New York 11794, USA
[6]These authors contributed equally

**Table S1.** SDM Primers

| | |
|---|---|
| SDE2 Δ395-427 | TTCACCTCTGTTGCA TCTGTCAGAGGACTG |
| SDE2 Δ428-451 (S428*) | CAG CAA GAC TCT TCT aaG TCA GAG GAC TGG CA |
| SDE2 K444A | CCG GCT TTA TTT GCC gcG CCT TTG AAA GGG AA |
| SDE2 K444 K447A | TTT GCC GCG CCT TTG gcA GGG AAG AAA AAA GA |
| SDE2 G417A | GA CTG AAA TGT GGG GcC ACT CTG CAG GAG CG |
| SDE2 G417A L419E | AAA TGT GGG GCC ACT gaG CAG GAG CGG GCA GC |
| SDE2 G412A L413E G417A | AA CTG ATG GCC CTT GcA gaG AAA TGT GGG GCC AC |
| SDE2 C415A G417A T418A | GCC CTT GGA CTG AAA gcT GGG GCC GCT CTG CA |
| SDE2 A417S A418V | GGA CTG AAA GCT GGG tCC GtT CTG CAG GAG CGG GC |
| SF3A3 Δ253-285 | TTCTCCTCCTGGGAG AGTACCAAAGGAAAG |
| SF3A3 Δ286-311 | GCCCAGAGACTATTC GACACTGAAAGGAAC |
| SF3A3 Δ253-311 | TTCTCCTCCTGGGAG GACACTGAAAGGAAC |

**Table S2.** Chemicals & Reagents

| Chemical/Reagent | Manufacturer | Cat. # |
| --- | --- | --- |
| 5-ethynyl-2´-deoxyuridine (EdU) | ThermoFisher Scientific | A10044 |
| Accugel 29:1 (40%) | National Diagnostics | EC-852 |
| Amicon Ultra Centrifugal Filter Devices, 3K | MilliporeSigma | UFC500308 |
| Amicon Ultra Centrifugal Filter Devices, 30K | MilliporeSigma | UFC503008 |
| Biotin azide | ThermoFisher Scientific | B10184 |
| BL21(DE3) Competent E. coli | New England Biolabs | C2527H |
| Click-iT® Cell Reaction Buffer Kit | ThermoFisher Scientific | C10269 |
| cOmplete, EDTA-free protease inhibitor cocktail | MilliporeSigma | 11873580001 |
| DL-Dithiothreitol (DTT) | MilliporeSigma | D0632 |
| Doxycycline hyclate | MilliporeSigma | D9891 |
| Duolink® In Situ Detection Reagents Red | MilliporeSigma | DUO92008 |
| Duolink® In Situ PLA® Probe Anti-Rabbit PLUS | MilliporeSigma | DUO92002 |
| Duolink® In Situ PLA® Probe Anti-Mouse MINUS | MilliporeSigma | DUO92004 |
| FLAG M2 affinity gel | MilliporeSigma | A2220 |
| Hen egg white lysozyme | MilliporeSigma | L4919 |
| Hexadimethrine bromide (polybrene) | MilliporeSigma | H9268 |
| HRV 3C protease (2 units/μL) | Pierce/ThermoFisher Scientific | 88946 |
| Isopropyl β-d-1-thiogalactopyranoside (IPTG) | MilliporeSigma | I6758 |
| GeneJuice Transfection Reagent | MilliporeSigma | 70967 |
| Glutathione agarose | Pierce/ThermoFisher Scientific | 16100 |
| GSTrap FF Column (1 x 5 mL) | Cytiva/GE | 17513101 |
| L-glutathione, reduced (GSH) | MilliporeSigma | G4251 |
| Phenylmethanesulfonyl fluoride (PMSF) | MilliporeSigma | P7626 |
| PES Protein Concentrator, 30K | Pierce/ThermoFisher Scientific | 88529 |
| Puromycin | MilliporeSigma | P8833 |
| RNAi MAX Transfection Reagent | ThermoFisher Scientific | 13778150 |
| Streptavidin magnetic beads | Pierce/ThermoFisher Scientific | 88816 |

| | | |
|---|---|---|
| Tris-borate-EDTA (TBE) buffer, 10X | National Diagnostics | EC-860 |
| Triton X-100 | MilliporeSigma | T8787 |
| Xfect Transfection Reagent | Clontech Laboratories | 631317 |

**Table S3.** Oligonucleotides

| | |
|---|---|
| EMSA1 (61 nt) | FAM-GACGCTGCCGAATTCTACCAGTGCCTTGCTAGGACATCTTTGCCCACCTGCAGGTTCACCC |
| EMSA2 (61 nt) (full complement) | GGGTGAACCTGCAGGTGGGCAAAGATGTCCTAGCAAGGCACTGGTAGAATTCGGCAGCGTC |
| EMSA3 (25 nt) | GCTGTCTAGAGGATCCGACTATCGA |
| EMSA4 (25 nt) | TGGGTGAACCTGCAGGTGGGCAAAGA |
| EMSA5 (62 nt) (half complement) | ATCGATAGTCGGATCCTCTAGACAGCTCCATGTAGCAAGGCACTGGTAGAATTCGGCAGCGT |
| EMSA6 (25 nt) | FAM-TCGATAGTCGGATCCTCTAGACAGC |
| DNA pulldown (80 nt) | BIO-$(TTT)_{10}$ CTCCCTTCTTCTCCTCCCTCTCCCTTCCC $(TTT)_7$ |
| DNA pulldown complement | $(AAA)_7$ GGGAAGGGAGAGGGAGGAGAAGAAGGGAG $(AAA)_{10}$ |
| NMR 16-mer | GCTATGGAGAACGGTA |
| IRE RNA (28 nt) | UCCUGCUUCAACAGUGCUUGGACGGAAC |

**Table S4.** NMR Spectrum Parameters

| Spectrum | Bruker Pulse Sequence | Size of FID (TD) | Spectral Width in ppm (SW) | NS | Others |
|---|---|---|---|---|---|
| N-HSQC | trosyf3gpphsi19.2 | F1-$^{1}$H 1024/ F2-$^{15}$N 128 | F1-$^{1}$H 12.00/ F2-$^{15}$N 36.00 | 16 | -- |
| HNCA | hncagp3d | F1-$^{1}$H 2048/ F2-$^{15}$N 40/ F3-$^{13}$C 64 | F1-$^{1}$H 15.94/ F2-$^{15}$N 35.00/ F3-$^{13}$C 30.00 | 16 | -- |
| HNCACB | hncacbgp3d | F1-$^{1}$H 2048/ F2-$^{15}$N 40/ F3-$^{13}$C 128 | F1-$^{1}$H 14.0029/ F2-$^{15}$N 35.00/ F3-$^{13}$C 80.00 | 16 | -- |
| HNCOCA | hncocagp3d | F1-$^{1}$H 2048/ F2-$^{15}$N 40/ F3-$^{13}$C 128 | F1-$^{1}$H 15.94/ F2-$^{15}$N 35.00/ F3-$^{13}$C 30.00 | 16 | -- |
| HNCACO | hncacogp3d | TD: F1-$^{1}$H 2048/ F2-$^{15}$N 40/ F3-$^{13}$C 128 | F1-$^{1}$H 14.0029/ F2-$^{15}$N 35.00/ F3-$^{13}$C 14.00 | 16 | -- |
| HNCOCACB | hncocacbgp3d | F1-$^{1}$H 2048/ F2-$^{15}$N 40/ F3-$^{13}$C 128 | F1-$^{1}$H 15.94/ F2-$^{15}$N 35.00/ F3-$^{13}$C 80.00 | 16 | -- |
| HNCO | hncogpwg3d | F1-$^{1}$H 2048/ F2-$^{15}$N 40/ F3-$^{13}$C 128 | F1-$^{1}$H 14.00/ F2-$^{15}$N 35.00/ F3-$^{13}$C 14.00 | 32 | -- |
| C-HSQC | hsqcetgp | F1-$^{1}$H 1024/ F2-$^{13}$C 256 | F1-$^{1}$H 15.94/ F2-$^{13}$C 165.0 | 16 | -- |
| N-TOCSY | dipsihsqcf3gpsi3d | F1-$^{1}$H 2048/ F2-$^{15}$N 40/ F3-$^{1}$H 64 | F1-$^{1}$H 15.94/ F2-$^{15}$N 35.00/ F3-$^{1}$H 15.94 | 16 | -- |
| CCCONH | ccconhgp3d | F1-$^{1}$H 2048/ F2-$^{15}$N 40/ F3-13C 128 | F1-$^{1}$H 14.00/ F2-$^{15}$N 35.00/ F3-$^{13}$C 80.00 | 16 | -- |
| C-NOESY | noesyhsqcetgp3d | F1-$^{1}$H 2048/ F2-$^{15}$N 40/ F3-$^{1}$H 128 | F1-$^{1}$H 14.00/ F2-$^{13}$C 75.00/ F3-$^{1}$H 14.00 | 32 | Mixing Time (D8): 0.2 sec |
| N-NOESY | noesyhsqcetf3gp3d | F1-$^{1}$H 2048/ F2-$^{15}$N 40/ F3-$^{1}$H 128 | F1-$^{1}$H 14.00/ F2-$^{15}$N 36.00/ F3-$^{1}$H 14.00 | 32 | Mixing Time (D8): 0.2 sec |
| IPAP | hsqcetf3gpiaphsisp | F1-$^{1}$H 1024/ F2-$^{15}$N 600 | F1-$^{1}$H 12.46/ F2-$^{15}$N 34.00 | 120 | -- |
| ssNOE | hsqcfpf3gpphwg | F1-$^{1}$H 2048/ F2-$^{15}$N 512 | F1-$^{1}$H 13.02/ F2-$^{15}$N 34.00 | 32 | Recovery Delay (D7): 7 sec |

F1, 2, 3 – Nucleus are the channels and its corresponding nucleus.

Supplementary Figure S1

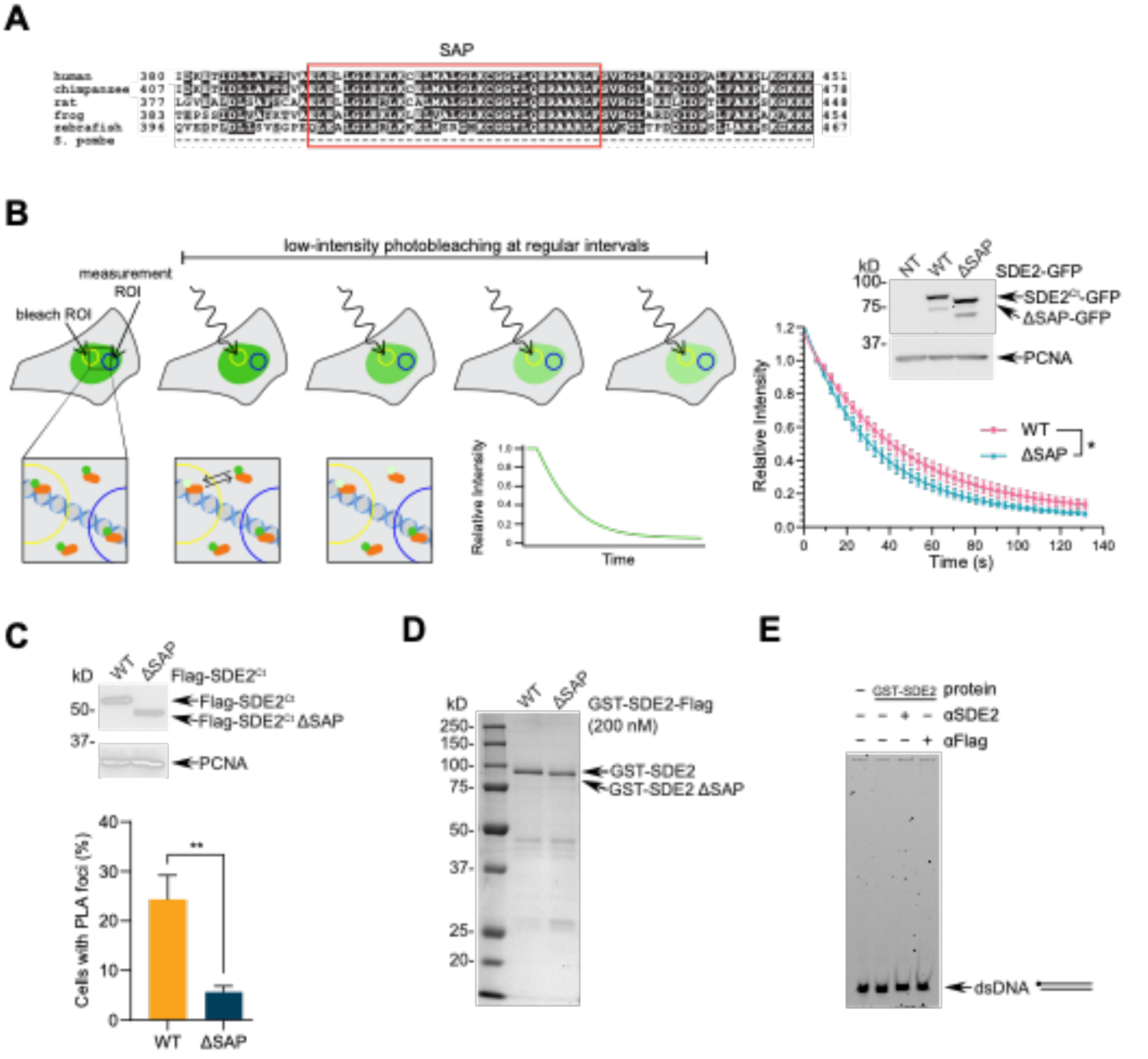


**Supplementary Figure S1. Characterization of DNA binding by SDE2$^{SAP}$**

**(A)** The sequence alignment of the SDE2 SAP domain from multiple species. The core SAP domain is marked with a red box. **(B)** Left: schematic showing how fluorescence loss in photobleaching (FLIP) is used to compare movement rates of nuclear proteins. Fluorophore-tagged proteins in the bleach region-of-interest (ROI) are partially photobleached and are allowed to exchange with non-photobleached proteins outside of the ROI, either through active movement or diffusion, before being bleached again. Bleaching steps are repeated at regular intervals until fluorescence signal is abolished or no longer decreases. Right: U2OS cells transiently expressing either SDE2-GFP WT or ΔSAP (Δ395-427) were subjected to FLIP while capturing images approximately every 3.5 s; curve fit to two-phase decay model, error bar: SD, *$p$<0.05, two-tailed t test. Western blots show proteins used for FLIP experiments. **(C)** Top: Western blot showing proteins used for SIRF experiments. U2OS cells were transfected with either Flag-SDE2$^{Ct}$ WT or ΔSAP and analyzed via Western blotting. Bottom: Percentage of replicating cells (positive for EdU-EdU signal) that contain PLA foci. Quantification of three independent experiments; Error bar: SEM, $^{**}p<0.01$, two-tailed Mann-Whitney test. **(D)** Coomassie Blue staining of the purified GST-SDE2-Flag WT and ΔSAP used for EMSAs. **(E)** Control EMSA experiment for supershift. Purified GST-SDE2-Flag were incubated with 60-mer dsDNA, additionally incubated with antibodies at a 1:3 ratio where indicated. DNA: 50 nM, protein: 200 nM. Lane 1: no protein control; lane 2: GST-SDE2-Flag; lane 3: GST-SDE2-Flag + anti-SDE2; lane 4: GST-SDE2-Flag + anti-Flag. Neither dsDNA binding nor supershift was observed.

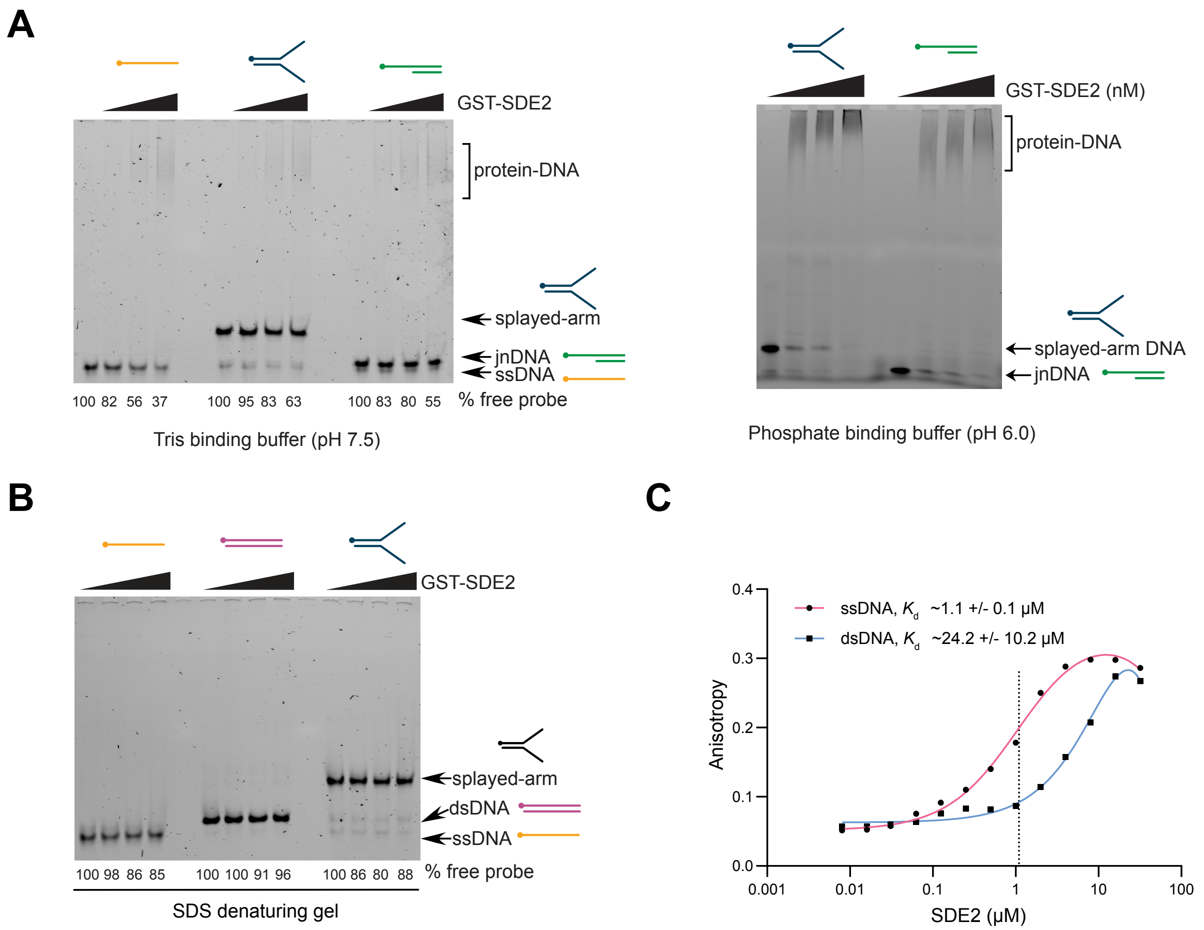


**Supplementary Figure S2. ssDNA-specific DNA binding of SDE2$^{SAP}$**

**(A)** EMSA of purified GST-SDE2-Flag incubated with 60-mer ssDNA, saDNA, or jnDNA (junction DNA). DNA: 50 nM, protein: 50-200 nM. Note that EMSAs were performed in both Tris binding (pH 7.5) and phosphate binding (pH 6.0) buffers, showing that results are reproducible in independent buffers. **(B)** EMSA of purified GST-SDE2-Flag incubated with 60-mer ssDNA, dsDNA, or saDNA then run on a *denaturing* gel. DNA: 50 nM, protein: 50-200 nM. **(C)** Increasing amounts of full-length GST-SDE2-Flag were incubated with FAM-labeled 60-mer ssDNA vs. dsDNA and analyzed using fluorescence anisotropy. Full-length SDE2 binds ssDNA with a dissociation constant ($K_d$) of approximately 1.1 µM, but to dsDNA with a $K_d$ of approximately 24 µM. Data fit to a one-site, total binding saturation curve. Dissociation constants are approximate due to challenges in reaching >90% purity for GST-SDE2-Flag at the scale required, due to its propensity to cleave in the large flexible region that makes up the bulk of the primary sequence.

#### Supplementary Figure S3

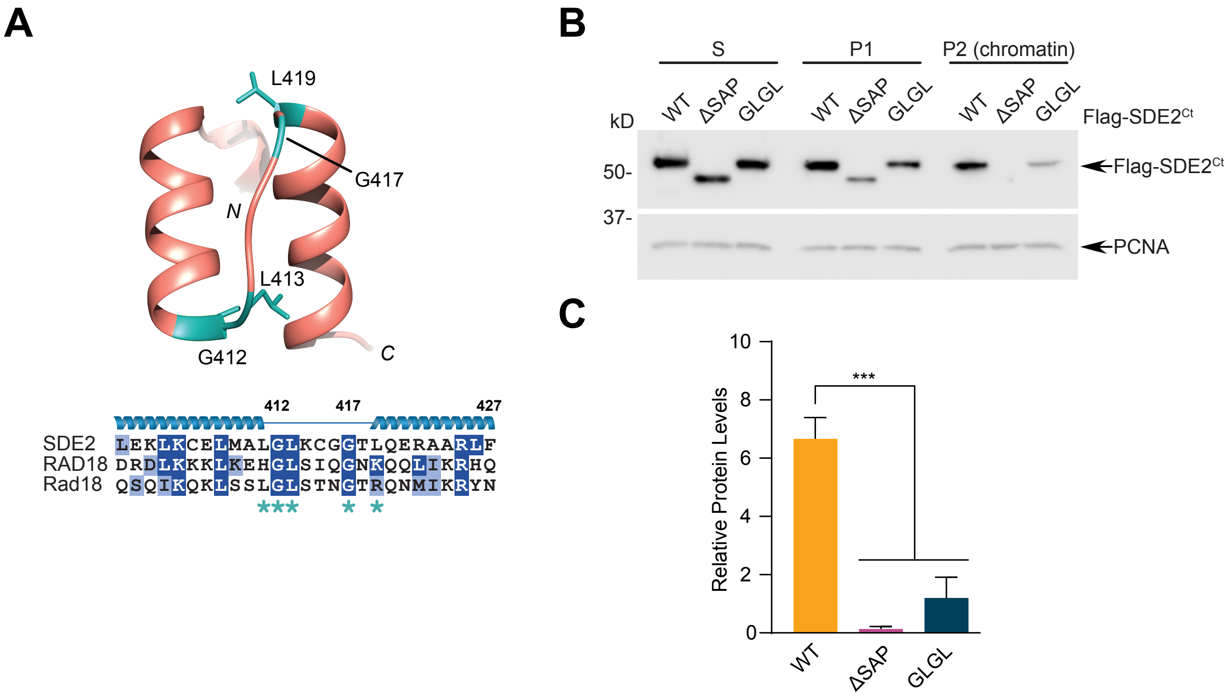


**Supplementary Figure S3. Conserved loop-region residues are important for DNA binding. (A)** A predicted structure of SDE2$^{SAP}$ (aa. 395-427), generated by Phyre2 based on primary sequence identity to known structures, with the most contributions coming from PDBs 2KVU (myocardin-like protein 1), 1ZRJ (E1B-55kDa-associated protein 5), 2DO1 (Hcc-1), 1H1J & 2WQG (Tho1), and 2RNN (Siz1). Residues corresponding to Rad18$^{SAP}$ binding mutants are highlighted in teal. **(B)** U2OS cells were transfected with Flag-SDE2$^{Ct}$ WT, ΔSAP (Δ395-427), or G412A/L413E/G417A/L419E (GLGL), fractionated to separate cytosolic and chromatin-associated protein pools, and analyzed by Western blotting. **(C)** Quantification of (B) from three independent experiments; error bar: SEM, ***$p$<0.001, one-way ANOVA.

A

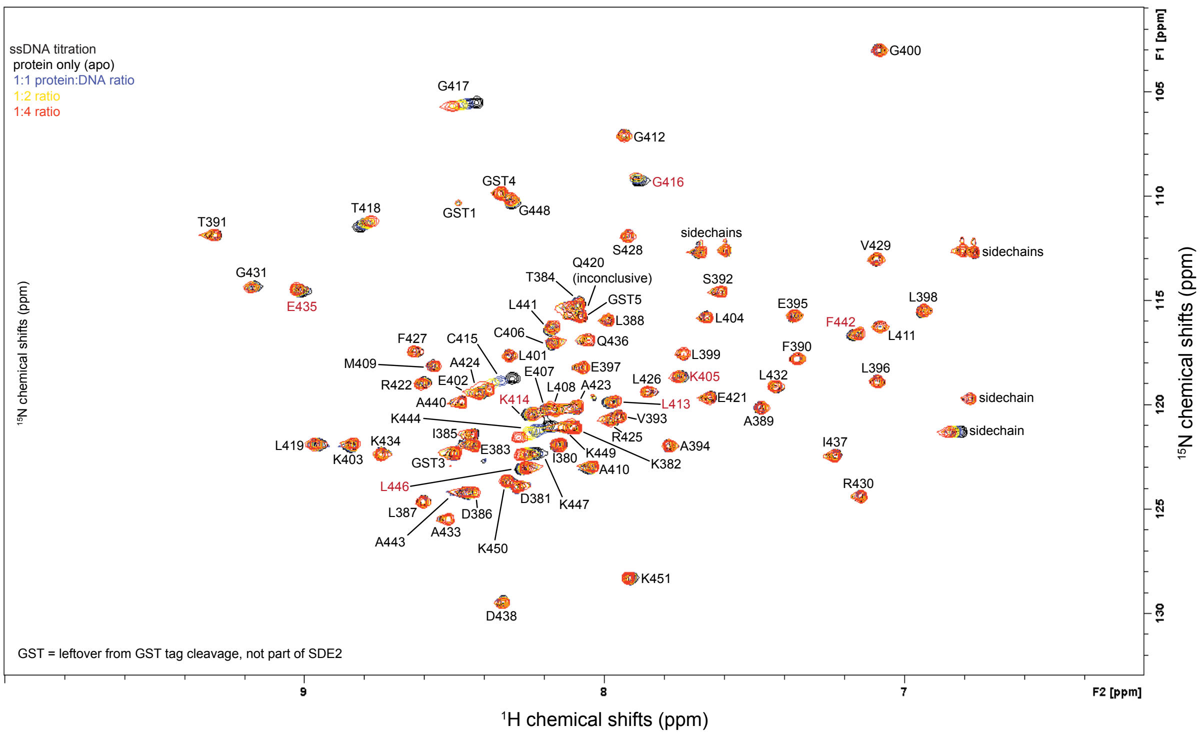


B

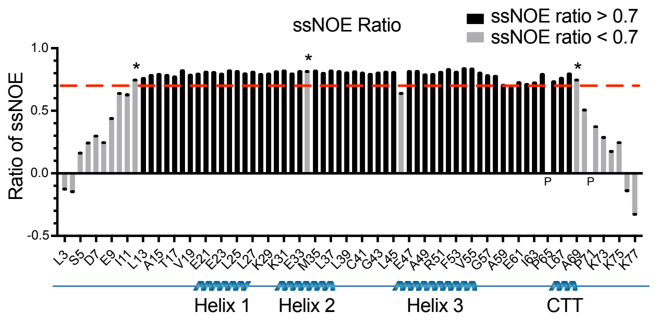


**Supplementary Figure S4. HSQC analysis of SDE2$^{SAP}$**

**(A)** Full assignment of $^{1}$H-$^{15}$N HSQC spectra, showing all cross-peak assignments. Cross-peaks 1-5 are from residues left after the cleavage of the GST tag by the HRV 3C protease, and are not part of the SAP+CTT motif (see Figure 3A). **(B)** Backbone dynamics of apo SDE2$^{SAP+CTT}$. Plot of {$^{1}$H}-$^{15}$N steady-state NOE (ssNOE) ratio vs. residue number. A ratio of >0.7 (indicated by red dashed line) was used as the cutoff for selection of the TALOS (Torsion Angle Likelihood Obtained from Shift and sequence similarity)-predicted dihedral angle and structural calculation. P indicates prolines, which do not show up in {$^{1}$H}-$^{15}$N HSQC, while * indicates residues with strongly overlapping cross-peaks for which ssNOE values could not be accurately extracted.

### Supplementary Figure S5

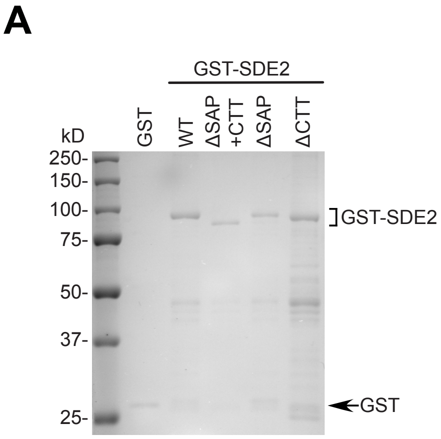


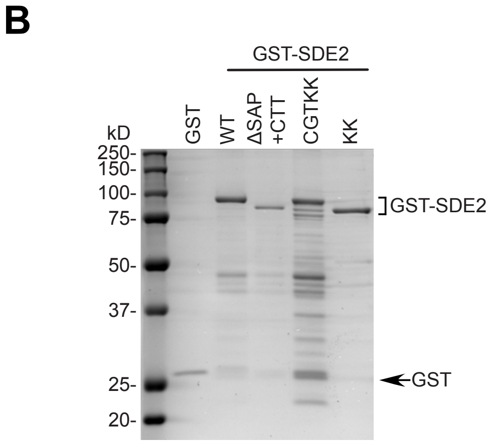


**Supplementary Figure S5. Contribution of SDE2$^{SAP}$ and SDE2$^{CTT}$ for DNA binding**
**(A)** Coomassie blue staining of purified GST-SDE2 variants used for EMSAs in Fig. 5D. **(B)** Coomassie blue staining of purified GST-SDE2 variants used for EMSAs in Figs. 5F and 5G.

### Supplementary Figure S6

A

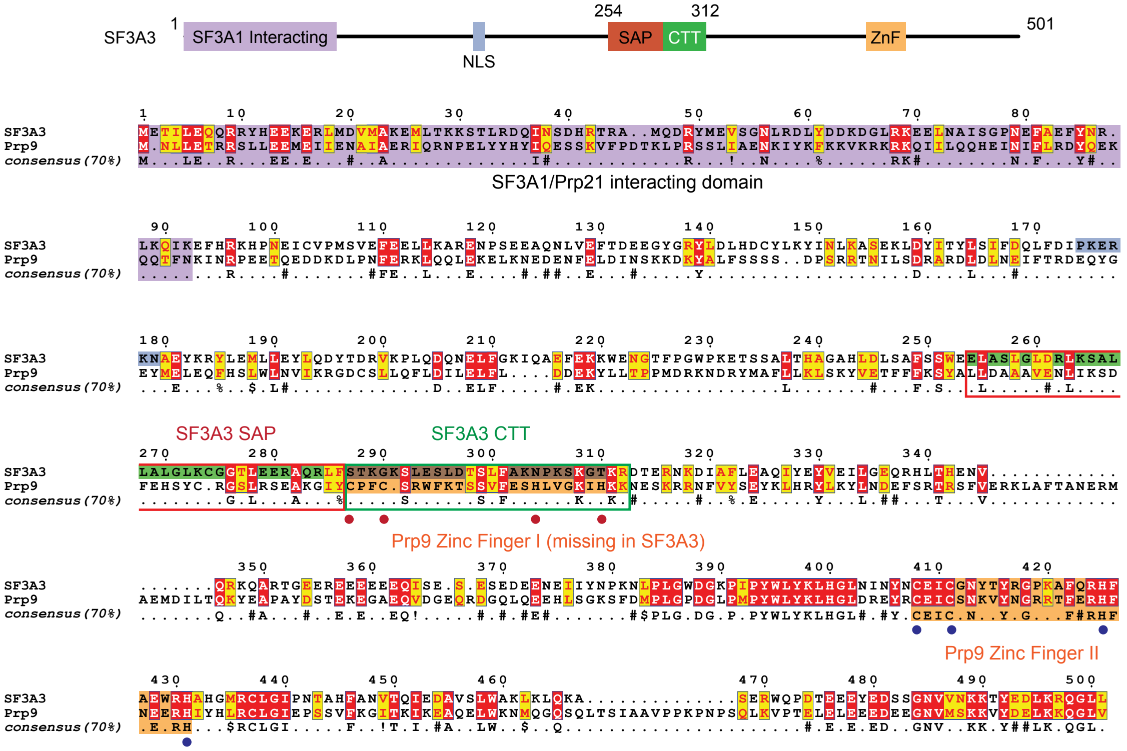


**Supplementary Figure S6. Conservation of the extended SAP in SDE2 and SF3A3**

**(A)** Sequence alignment of SF3A3 and its yeast homolog Prp9. The SF3A1/Prp21 interacting domain is colored purple, SF3A3 SAP and CTT are colored red and green, respectively, and the zinc fingers are colored orange. Notice that while SF3A3 and Prp9 both share the C-terminal ZnF motif that binds the spliceosome 15S intermediate, Prp9 contains an additional zinc finger at the location of the CTT in SF3A3. The positions of missing cysteine and histidine residues in the ZnF motif I of SF3A3 are marked with red dots.

### Supplementary Figure S7

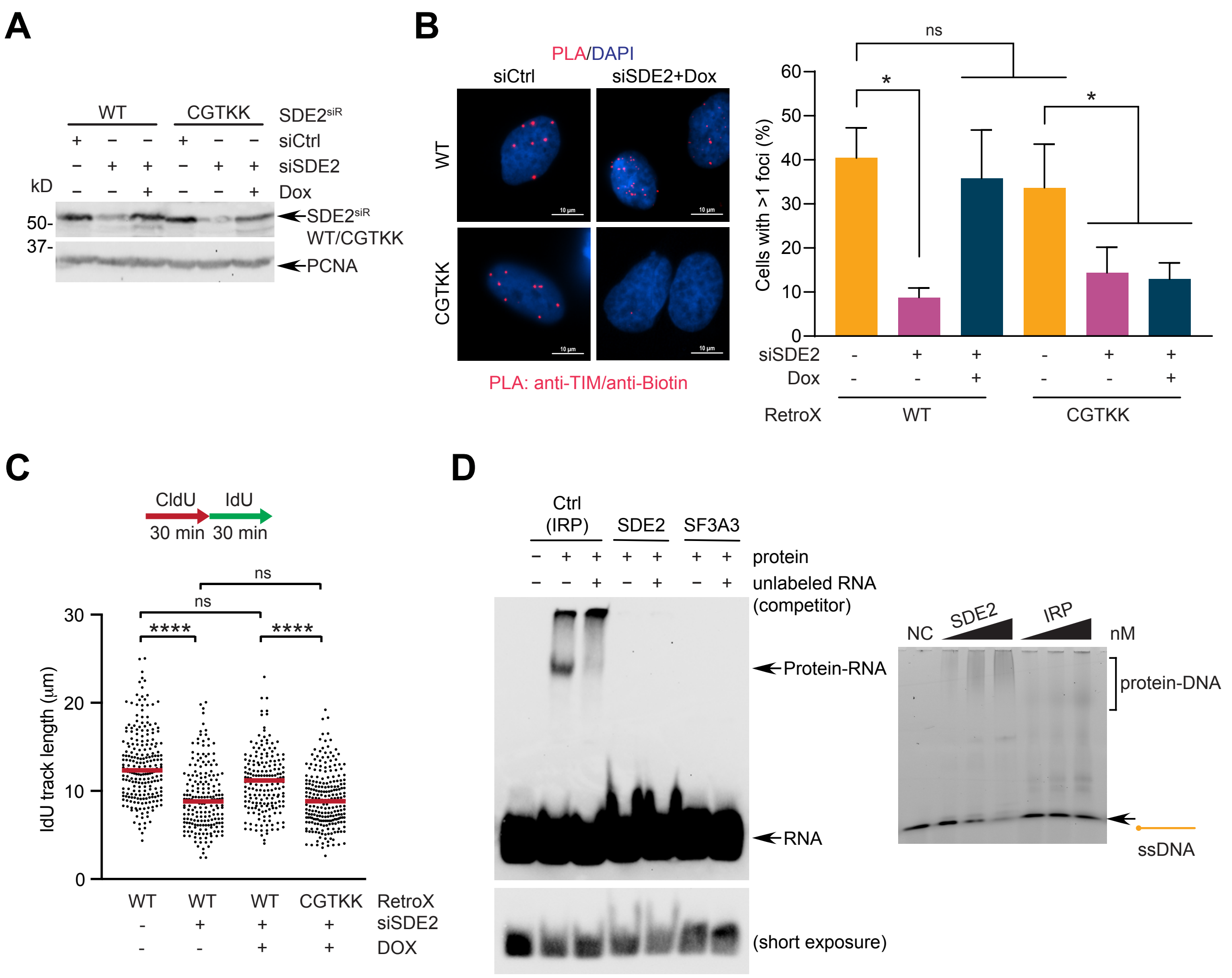


**Supplementary Figure S7. Characterization of the SAP point mutants**

**(A)** Western blotting that shows the complementation of SDE2 WT or SAP mutant (CGTKK; C415A/G417S/T418V/K444A/K447A) in Retro-X cells by siRNA transfection and Dox induction. **(B)** Left: Representative images of TIM:EdU PLA foci in the Retro-X SDE2 WT or SAP mutant cells following SDE2 siRNA transfection and dox induction. Scale bar: 10 µm. Right: quantification of cells positive for TIM-EdU PLA foci normalized to EdU-EdU PLA-positive replicating cells (>300 cells per condition, n=3, error=SEM, *$p<0.05$, 2-way ANOVA). **(C)** Dot plots of the DNA fiber IdU track length from Retro-X cells reconstituted with SDE2 WT or CGTKK point mutant (>200 tracks per condition, ****$p<0.0001$, ns, not significant, Mann-Whitney). **(D)** Left: competitive RNA electrophoretic mobility shift assay (REMSA) of control protein (iron-responsive protein: IRP), purified GST-SDE2, or GST-SF3A3 incubated with 28-mer 3' biotin-labeled IRE (iron-responsive element) RNA. Competitor is unlabeled IRE RNA and shows that binding is specific. biotin-RNA: 6.25 nM, unlabeled RNA 1 µM. Right: EMSA of GST-SDE2 or IRP incubated with 60-mer ssDNA, showing that IRP does not bind to ssDNA.
